# G protein-regulated endocytic trafficking of adenylyl cyclase type 9

**DOI:** 10.1101/2020.04.23.057026

**Authors:** André M. Lazar, Roshanak Irannejad, Tanya A. Baldwin, Aparna A. Sundaram, J. Silvio Gutkind, Asuka lnoue, Carmen W. Dessauer, Mark von Zastrow

## Abstract

GPCRs are increasingly recognized to initiate signaling via heterotrimeric G proteins as they move through the endocytic network, but little is known about how relevant G protein effectors are localized. Here we report dynamic trafficking of adenylyl cyclase type 9 (AC9) from the plasma membrane to endosomes, while adenylyl cyclase type 1 (AC1) remains in the plasma membrane, and stimulation of AC9 trafficking by ligand-induced activation of Gs-coupled GPCRs or Gs. AC9 transits a similar dynamin-dependent early endocytic pathway as activated GPCRs but, in contrast to GPCR trafficking which is regulated by β-arrestin but not Gs, AC9 trafficking is regulated by Gs but not β-arrestin. We also show that AC9, but not AC1, contributes to cAMP production from endosomes. These results reveal dynamic and isoform-specific trafficking of adenylyl cyclase in the endocytic network, and a discrete role of a heterotrimeric G protein in controlling subcellular location of a relevant effector.

## Introduction

G protein-coupled receptors (GPCRs) comprise the largest family of signaling receptors and an important class of therapeutic drug targets (Lefkowitz, 2007; Rosenbaum et al., 2009). GPCRs are so-named because a major mechanism by which they mediate transmembrane signaling is through ligand-dependent activation of heterotrimeric G proteins that act as intracellular signal transducers (Gilman, 1987; Hilger et al., 2018; Spiegel, 1987; Sunahara, 1996). Canonical GPCR signaling invariably requires one additional component, an effector protein which is regulated by the G protein to convey the signal downstream (Dessauer et al., 1996; Gilman, 1987; Rosenbaum et al., 2009). Ligand-dependent signaling by GPCR - G protein - effector cascades was thought for many years to be restricted to the plasma membrane, with endocytosis considered only in the context of signal termination and homeostatic down-regulation of receptors (Di Fiore and von Zastrow, 2014; Lohse and Calebiro, 2013; Lohse and Hofmann, 2015). This view has expanded over the past several years, with accumulating evidence that GPCR and G protein activation by ligands can also occur internally.

Signaling mediated by the beta-2 adrenergic receptor (β2AR) provides a clear example, and the β2AR is generally considered a model for the GPCR family more broadly (Lefkowitz, 2007; Rosenbaum et al., 2009). The β2AR initiates signaling primarily by coupling to the stimulatory heterotrimeric G protein, Gs, at the plasma membrane. β2ARs then internalize by ligand-dependent accumulation into clathrin-coated pits and continuously cycle between the plasma membrane and endosomes (von Zastrow and Kobilka, 1994, 1992). This process is initiated by receptor phosphorylation and binding to β-arrestins at the plasma membrane (Ferguson et al., 1996; Goodman et al., 1996), events which were shown previously to prevent β2AR coupling to Gs (Lohse et al., 1990). This overall sequence, recognized even before its mechanistic underpinnings were elucidated (Harden et al., 1980), supported a long-held view that β2ARs are unable to engage G proteins once internalized. This view changed with the finding that β2ARs initiate a second phase of ligand-dependent activation of Gs shortly after arriving in the limiting membrane of early endosomes (Irannejad et al., 2013). Numerous GPCRs have now been shown or proposed to activate Gs after endocytosis, and to generate a discrete ‘wave’ of signaling internally (Irannejad et al., 2015; Lohse and Calebiro, 2013; Thomsen et al., 2016; Vilardaga et al., 2014).

Gs transduces downstream effects by stimulating adenylyl cyclases (ACs) to produce cyclic AMP (cAMP). cAMP is an extensively studied diffusible second messenger (Lohse and Hofmann, 2015; Sutherland, 1971; Taylor et al., 2013), and studies of the cAMP system provided much of the early evidence suggesting that GPCRs have the capacity to activate G proteins after they internalize (Calebiro et al., 2009; Clark et al., 1985; Ferrandon et al., 2009; Kotowski et al., 2011; Mullershausen et al., 2009; Slessareva et al., 2006). Nine transmembrane AC isoforms are conserved in mammals, each stimulated by Gs but differing in regulation by other G proteins and signaling intermediates, and cells typically coexpress more than one isoform (Sadana et al., 2009; Sunahara, 1996). Biochemical and structural aspects of regulated cAMP production by ACs have been extensively studied, but much less is known about the cellular biology of ACs.

According to the present understanding, GPCR-stimulated cAMP production requires core components of the signaling cascade - the GPCR, Gs and AC - to be in the same membrane bilayer (Gilman, 1989). Whereas GPCRs and Gs are well known to undergo dynamic redistribution between the plasma membrane and endocytic membranes (Allen et al., 2005; Hynes et al., 2004; Irannejad et al., 2015; Marrari et al., 2007; von Zastrow and Kobilka, 1992; Wedegaertner et al., 1996), much less is known about the subcellular organization or dynamics of transmembrane ACs. Nevertheless, adenylyl cyclase activity was noted on intracellular membranes many years ago (Cheng and Farquhar, 1976), and recent functional evidence has implicated several AC isoforms in GPCR-regulated cAMP production from endomembrane compartments (Caldieri and Sigismund, 2016; Cancino et al., 2014; Ferrandon et al., 2009; Vilardaga et al., 2014). Critical gaps in current knowledge are how ACs localize to relevant internal membranes, if the subcellular localization of ACs is selective among isoforms, and if it is regulated. We make initial inroads into this frontier by delineating the dynamic and isoform-specific endocytic trafficking of AC9 and its ligand-dependent regulation by Gs.

## Results

### Regulated and isoform-selective trafficking of AC9 to endosomes

Human embryonic kidney (HEK293) cells comprise a well established model system for investigating GPCR signaling via the cAMP cascade, particularly signaling initiated by β2ARs which are endogenously expressed in these cells (Violin et al., 2008). The β2AR stimulates cAMP production primarily from the plasma membrane, with endosomal activation contributing a relatively small but functionally significant fraction (Irannejad et al., 2013; Tsvetanova and von Zastrow, 2014). In a similar vein, AC3 and AC6 are the most highly expressed AC isoforms and mediate most of global cAMP production in these cells, but AC1 and AC9 transcripts are expressed at somewhat lower levels and comparable to one another (Soto-Velasquez et al., 2018). This prompted us to ask whether AC1 and/or AC9 might be relevant to generating the portion of cellular cAMP initiated by β2AR activation in endosomes.

As an initial step to investigate this hypothesis, we examined the subcellular localization of AC1 and AC9 using a recombinant epitope tagging strategy, beginning with AC1 because this isoform was previously shown to tolerate an N-terminal Flag tag (Chen et al., 1997). When expressed in HEK293 cells, Flag-tagged AC1 (Flag-AC1) localized primarily to the plasma membrane (**Fig 1A**), similar to a co-expressed HA-tagged β2AR construct (HA-β2AR) (**Fig 1A**). Application of the β-adrenergic agonist isoproterenol caused HA-β2ARs to redistribute within minutes to cytoplasmic punctae (**Fig 1A**), as described previously, but Flag-AC1 remained at the plasma membrane (**Fig 1A**). We confirmed this physical separation across hundreds of cotransfected cells (**Fig S1D**) and quantified it in two ways. First, we determined the number of internal punctae per cell in which HA-β2AR and Flag-AC1 colocalized. Second, we determined the fraction of cells visualized in each microscopic field that contained at least 10 such punctae. Both metrics verified selective internalization of the β2AR but not of AC1 (**Fig 1C** and **D**, left set of bars).

**Figure 1:**
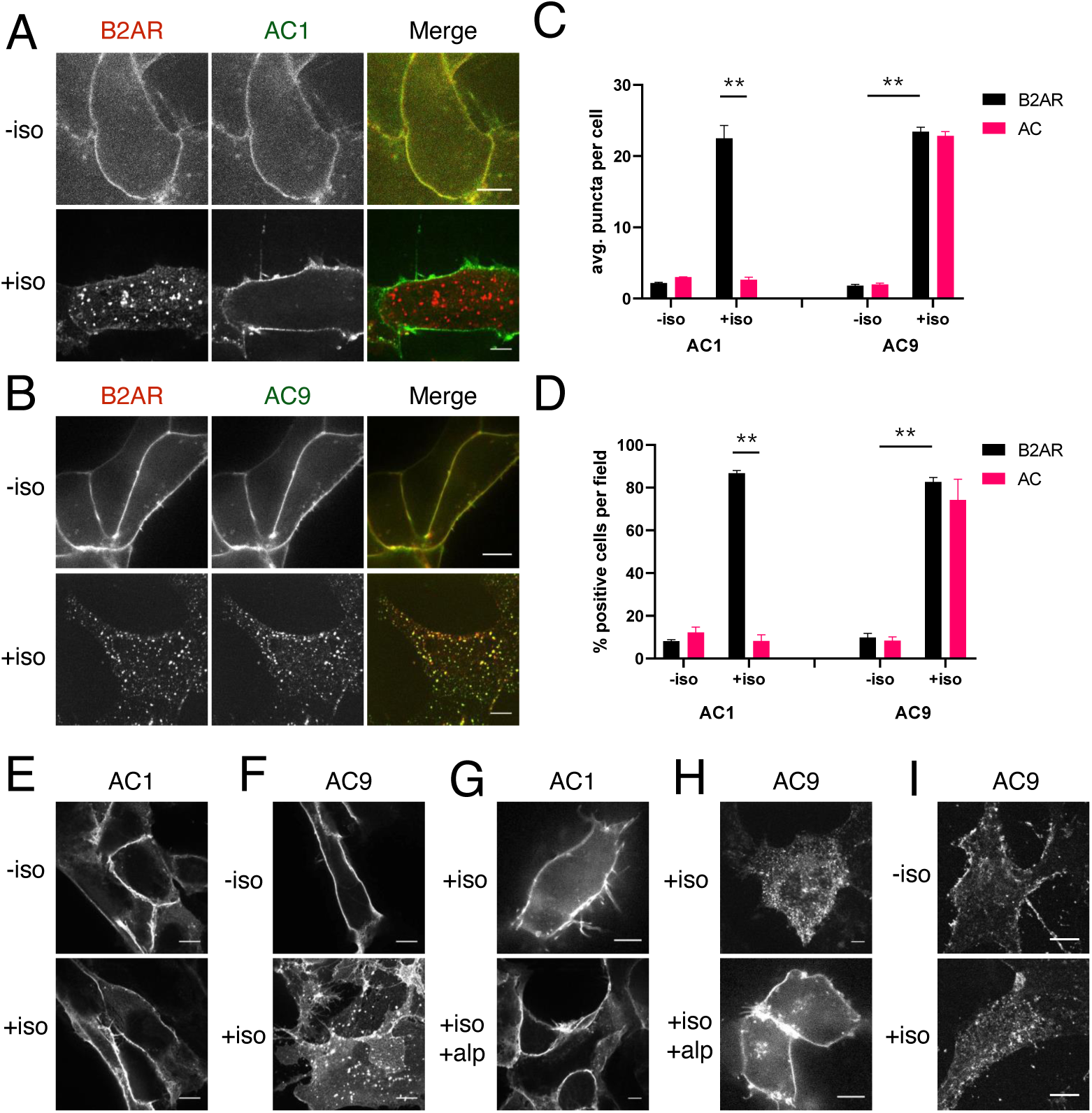
β2AR activation causes redistribution of AC9 but not AC1. **(A)** Representative confocal imaging of HEK293 cells coexpressing HA-β2AR and Flag-AC1 after treatment with 10 µM isoproterenol or control for 30 min. Scale Bar is 8 µm. **(B)** Representative confocal images of HEK293 cells coexpressing HA-β2AR and Flag-AC9 after treatment with 10 µM isoproterenol or control for 30 min. Scale Bar is 8 µm. **(C)** Quantification of internal puncta that are β2AR or AC1/9 positive, taken from wide field images (see: Supp. Fig 1D,1E) [mean±SEM; n=3 experiments, 10 visual fields and 200+ cells per condition]. ** P < 0.01 by two-tailed t-test. **(D)** Quantification of cells with >10 internal puncta that are β2AR or AC1/9 positive, taken from wide field images (see: Supp. Fig 1D,1E) [mean±SEM; n=3 experiments, 10 visual fields and 200+ cells per condition]. ** P < 0.01 by two-tailed t-test. **(E-F)** Representative confocal imaging of HEK293 cells expressing Flag-AC1 **(E)** or Flag-AC9 **(F)** after treatment with 10 µM isoproterenol or control for 30 min. Scale Bar is 8 µM. **(G-H)** Representative confocal imaging of HEK293 cells expressing Flag-AC1 **(G)** or Flag-AC9 **(H)**. Cells were stimulated with 100 nM isoproterenol for 30 min with or without 15 min of pretreatment with 10 µM alprenolol. **(l)** Representative confocal images of primary culture human airway smooth muscle cells immunostained for endogenous AC9 after treatment with 10 µM isoproterenol or control for 30 min. Scale Bar is 16 µm.

We applied a similar tagging strategy to AC9, and verified that Flag tagging also does not disrupt the functional activity of AC9 (**Fig S1A, S1B**). Flag-AC9 localized predominantly in the plasma membrane in the absence of agonist, similar to Flag-AC1 (**Fig 1C, 1D**). However, application of isoproterenol markedly increased Flag-AC9 localization to intracellular punctae, and the majority of AC9-containing punctae also contained internalized HA-β2AR (**Fig 1B**). Quantitative analysis (**Fig S1D**) verified accumulation of both β2AR and AC9 in the same endosomes (**Fig 1C** and **D**, left set of bars). We observed a similar localization of AC9 labeled in its C-terminal cytoplasmic domain with GFP (AC9-GFP), enabling live-cell confocal imaging which revealed rapid movement of AC9-containing endodomes (Supplemental Video 1). These results indicate that AC9 dynamically traffics to endosomes containing β2ARs, that this trafficking is isoform-specific because AC1 remains in the plasma membrane, and that it is regulated because endosomal accumulation of AC9 is increased by β2AR activation.

Isoproterenol also increased endosomal localization of Flag-AC9 in the absence of recombinant β2AR overexpression (**Fig 1E, 1F, S1C**), and this effect was blocked by the -adrenergic antagonist alprenolol (**Fig 1G, 1H, S1C**). This indicates that activation of endogenous β2ARs is sufficient to stimulate AC9 trafficking, and that AC9 trafficking is not an off-target drug effect. AC9, and ACs in general, are low-abundance proteins in HEK293 cells, and we were unable to reliably detect endogenous AC9 in HEK293 cells using available antibodies. However, we detected endogenous AC9 in primary human airway smooth muscle cells which express this protein at a higher level and also express β2ARs (Billington et al., 1999). Endogenous AC9 immunoreactivity localized primarily to the plasma membrane under basal conditions in these cells, and its localization to internal punctae increased after isoproterenol application (**Fig 1l, S1C**). This suggests that the trafficking behavior revealed by study of recombinant, tagged AC9 is relevant to the native protein.

### AC9 traffics via a similar pathway as β2AR but is differentially regulated

Endosomes that accumulate agonist-internalized β2ARs are marked by Early Endosome Antigen 1 (EEA1), and agonist-dependent coupling of the β2AR to Gs has been explicitly shown to occur in EEA1-associated endosomes (Irannejad et al., 2013). We established isoform-specific localization of Flag-AC9 in EEA1-marked endosomes by confocal microscopy (**Fig 2A, 2B**) and then applied a previously established endosome immunoisolation procedure, based on anti-EEA1 pulldown (Cottrell et al., 2009; Hammond et al., 2010; Temkin et al., 2011), to isolate these endosomes and probe their membrane composition biochemically. Both HA-β2AR and Flag-AC9 were enriched in the endosome fraction isolated from isoproterenol-treated cells co-expressing these constructs, as detected by immunoblot analysis (**Fig 2C**) and quantified over multiple experiments (**Fig 2D**). In contrast, HA-β2AR but not Flag-AC1 was enriched in parallel isolations from cells co-expressing HA-β2AR and Flag-AC1, with similar levels of overall protein expression across both cell populations verified in cell lysates (**Fig 2C, 2D**). Documenting separation efficiency and fraction purity, the endosome fraction recovered ∼34% of total cellular EEA1 but <5% of Golgi, endoplasmic reticulum, or plasma membrane markers (**Fig S2A**).

**Figure 2:**
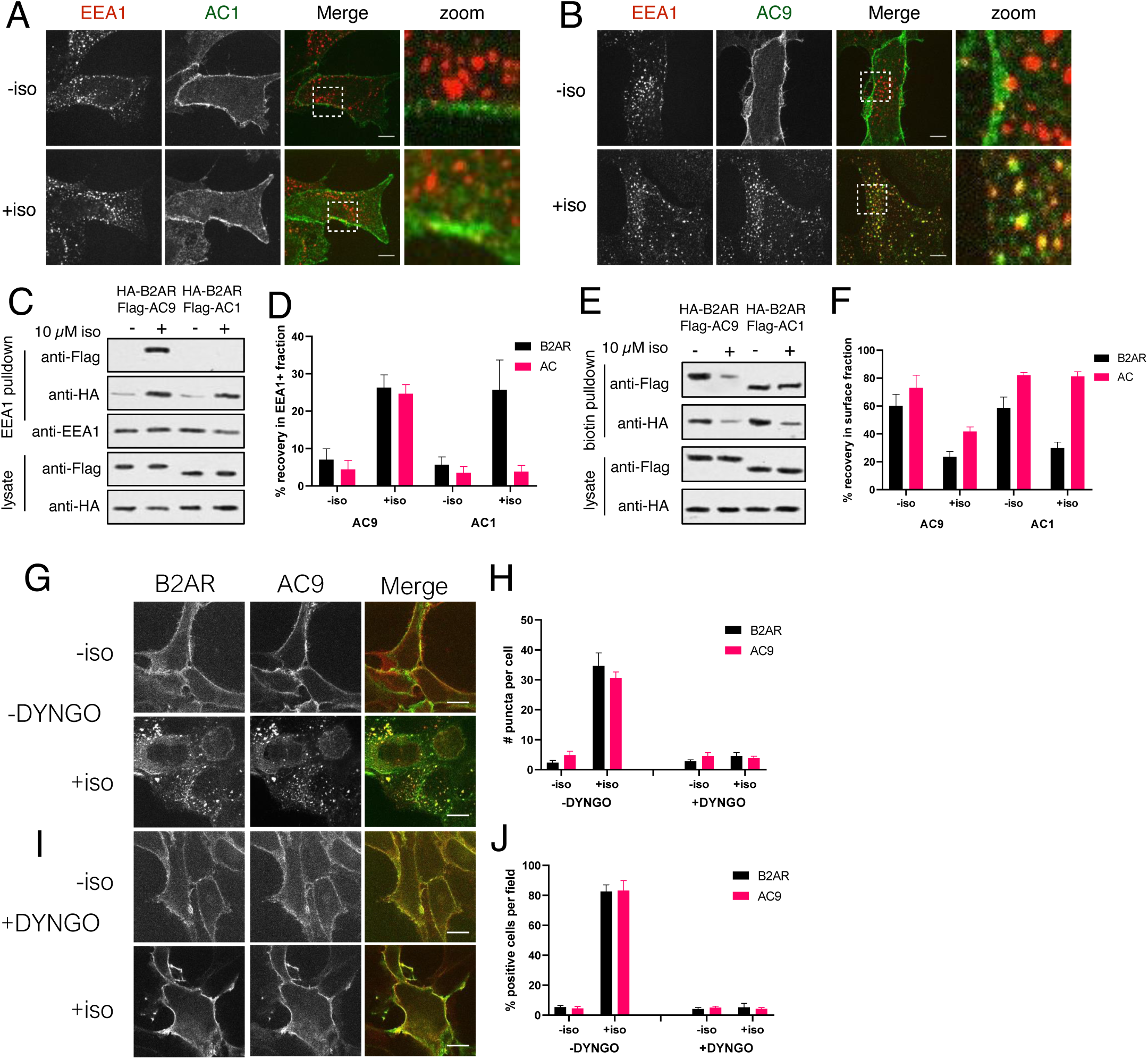
Surface AC9 is internalized to early endosomes upon adrenergic stimulation. **(A-B)** Representative confocal images of HEK293 cells expressing Flag-AC1 **(A)** or Flag-AC9 **(B)** after treatment with 10 µM isoproterenol or control for 30 min and stained for endogenous EEA1. Scale bar is 8 µm. **(C)** Representative western blot of a fraction isolated using antibodies to EEA1. Lanes 1-2 correspond to control HEK293 cells, lanes 3-4 to cells coexpressing Flag-AC9 and HA-β2AR, and lanes 5-6 to cells coexpressing Flag-AC1 and HA-β2AR. **(D)** Quantification of recovery of HA-β2AR, Flag-AC9 and Flag-AC1 in the endosome fraction relative to cell lysate. [mean±SEM; n=7 experiments]. ** P < 0.01 by two-tailed t-test. **(E)** Representative western blot of the surface exposed fraction isolated by surface labeling with Sulfo-NHS-biotin and purified with streptavidin. Lanes 1-2 correspond to cells coexpressing Flag-AC9 and HA-β2AR, and lanes 3-4 to cells coexpressing Flag-AC1 and HA-β2AR. **(F)** Quantification of recovery of HA-β2AR, Flag-AC9 and Flag-AC1 in the surface biotinylated fraction relative to total cell lysate. [mean±SEM; n=7 experiments]. ** P < 0.01 by two-tailed t-test. **(G)** Representative confocal images of HEK293 cells coexpressing HA-β2AR and Flag-AC9 after treatment with 10 µM isoproterenol or control for 30 min. Cells were treated with DMSO for 15 min prior to agonist exposure. **(H)** Quantification of internal puncta that are β2AR or AC9 positive, taken from wide field images (see: Supp. Fig 2D,2E) [mean±SEM; n=3 experiments, 10 visual fields and 200+ cells per condition]. ** P < 0.01 by two-tailed t-test. **(l)** Representative confocal images of HEK293 cells coexpressing HA-β2AR and Flag-AC9 after treatment with 10 µM isoproterenol or control for 30 min. Cells were treated with DYNGO-4a for 15 min prior to agonist exposure. **(J)** Quantification of cells with >10 internal puncta that are β2AR or AC1/9 positive, taken from wide field images (see: Supp. Fig 2D,2E) [mean±SEM; n=3 experiments, 10 visual fields and 200+ cells per condition]. ** P < 0.01 by two-tailed t-test.

As an additional biochemical verification, we used cell surface biotinylation to assess protein depletion from the plasma membrane (Flesch et al., 1995; Whistler and von Zastrow, 1998). Isoproterenol produced a marked reduction of Flag-AC9 in the surface-biotinylated fraction, but Flag-AC1 was unchanged (**Fig 2E, 2F**) and surface HA-β2AR decreased after isoproterenol application irrespective of whether AC1 or AC9 was coexpressed (**Fig 2E, 2F**). These observations independently demonstrate isoform-specific trafficking of AC9 from the plasma membrane regulated coordinately with trafficking of the β2AR.

A characteristic property of the clathrin-mediated endocytic pathway utilized by the β2AR is that it requires dynamin, an endocytic GTPase which can be acutely blocked by the chemical inhibitor DYNGO-4a (Irannejad et al., 2013). Whereas both Flag-β2AR and AC9-GFP accumulated in endosomes in the vehicle (0.1% DMSO) control condition (**Fig 2G, 2H**), endosomal accumulation of both proteins was blocked in the presence of DYNGO-4a (**Fig 2l, 2J**). These results suggest that AC9 trafficking to endosomes is mediated by a similar dynamin-dependent endocytic mechanism and pathway that is known to mediate regulated trafficking of the β2AR, as well as many other GPCRs.

Despite AC9 and β2AR trafficking via the same early endocytic pathway, they do so independently. An early clue to this was that AC9 trafficking can be reduced by pre-exposure of cells outside of the tissue culture incubator for a short period of time, whereas β2AR trafficking was relatively insensitive to this manipulation (see Methods). The molecular basis of this differential environmental sensitivity of AC9 relative to β2AR trafficking remains unknown, but it motivated us to examine in more detail mechanistic aspects of AC9 traffic control.

### AC9 trafficking is stimulated by Gs-but not Gi -coupled GPCRs

We first asked if the β2AR is unique in its ability to stimulate endosomal accumulation of AC9, or if AC9 trafficking can be stimulated by another Gs-coupled GPCR. To do so we focused on the vasopressin-2 receptor (V2R), a distinct Gs-coupled GPCR which also undergoes agonist-induced trafficking to early endosomes but is not endogenously expressed in HEK293 cells (Birnbaumer, 2000). As expected, the V2R agonist arginine-vasopressin (AVP) had no effect in cells not coexpressing HA-Vβ2AR, and Flag-AC9 remained in the plasma membrane irrespective of exposure to AVP (**Fig S3A**). In cells co-transfected with HA-V2R, however, AVP stimulated Flag-AC9 to accumulate in endosomes that also accumulated internalized Flag-V2R (**Fig 3A, 3D, 3E, S3D**), with internalization of both proteins confirmed by surface biotinylation (**Fig 3F, 3G**). These results indicate that the ability of GPCR activation to promote trafficking of AC9 to endosomes is not unique to the β2AR. Rather, it appears to be a more widespread property of GPCRs which couple to Gs.

**Figure 3:**
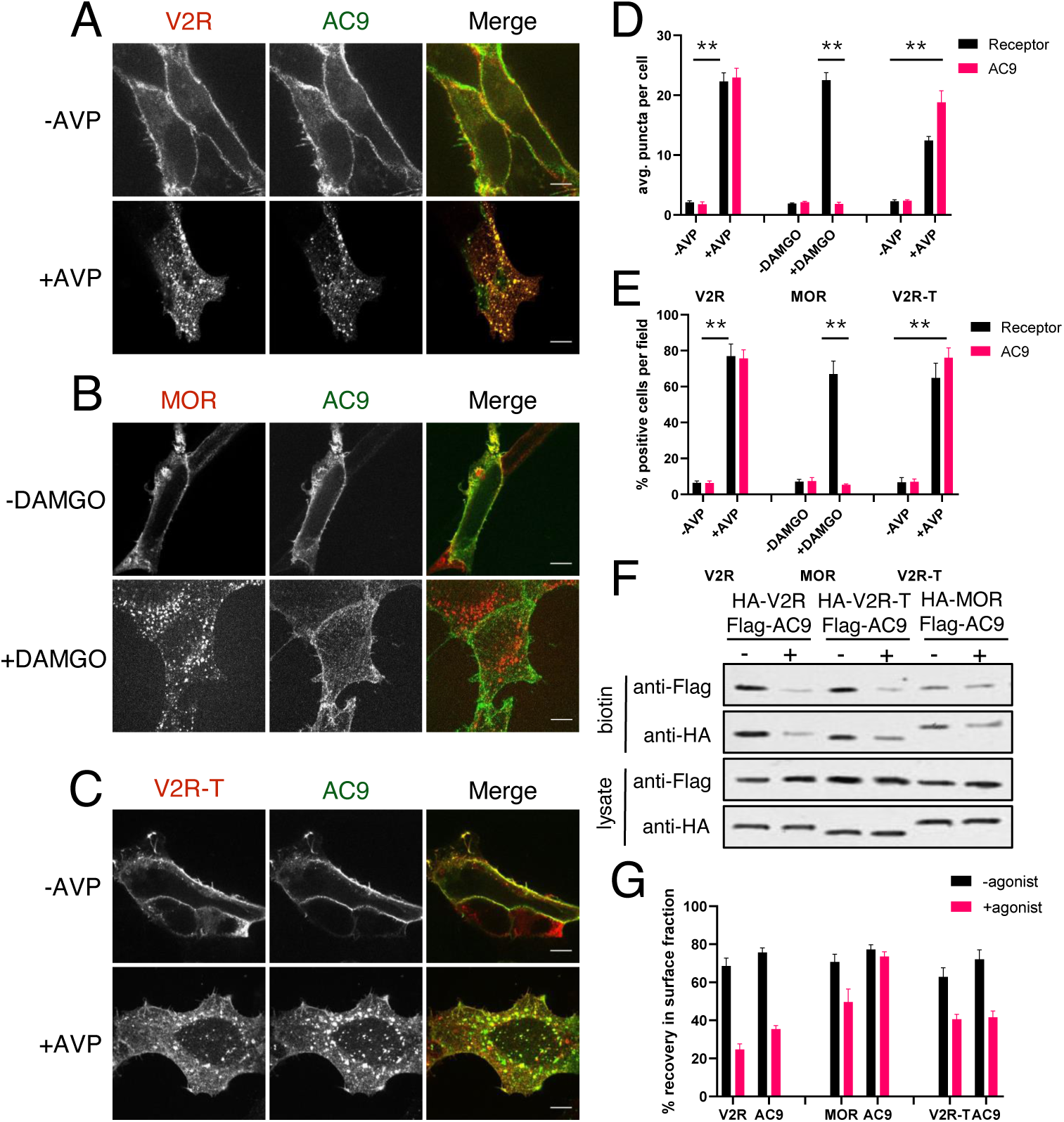
V2R, but not MOR, can cause AC9 to localize to early endosomes. **(A-C)** Representative confocal imaging of HEK293 cells coexpressing Flag-AC9 and HA-V2R **(A)**, HA-MOR **(B)**, or HA-V2R-trunc **(C)**, after treatment with 10 µM agonist (AVP or DAMGO) or control for 30 min. Scale Bar is 8 µm. **(D)** Quantification of internal puncta that are V2R, MOR, V2R-T, or AC1/9 positive, taken from wide field images (see: Supp. Fig 3D, 3E, 3F) [mean±SEM; n=3 experiments, 10 visual fields and 200+ cells per condition]. ** P < 0.01 by two-tailed t-test. **(E)** Quantification of cells with >10 internal puncta that are V2R, MOR, V2R-T or AC1/9 positive, taken from wide field images (see: Supp. Fig 3D, 3E, 3F) [mean±SEM; n=3 experiments, 10 visual fields and 200+ cells per condition]. ** P < 0.01 by two-tailed t-test. **(F)** Representative western blot of the surface biotinylated fraction from HEK293 cells coexpressing HA-V2R and Flag-AC9 (lanes 1-2), HA-V2R-T and Flag-AC9 (lanes 3-4), or HA-MOR and Flag-AC9 (lanes 5-6). **(G)** Recovery of tagged protein in the surface biotinylated fraction relative to the total cell lysate as seen in **(F)** [mean±SEM; n=7 experiments]. ** P < 0.01 by two-tailed t-test.

We next asked if the ability to stimulate AC9 trafficking extends to GPCRs that couple to other heterotrimeric G proteins. We focused on the µ-opioid receptor (MOP-R or MOR) because this GPCR transits the same early endocytic pathway (Keith et al., 1998) but couples to Gi, rather than Gs (Kieffer and Evans, 2009). As expected, application of the µ-opioid agonist [D-Ala^2^, N-MePhe^4^, Gly-ol]-enkephalin (DAMGO) stimulated transfected HA-MOR to accumulate in endosomes, but Flag-AC9 remained in the plasma membrane (**Fig 3B, 3D, 3E, S3E**). Selective internalization of HA-MOR but not Flag-AC9 was verified by surface biotinylation (**Fig 3F, 3G**). These results suggest that AC9 trafficking can be stimulated by GPCRs that couple to Gs but not Gi.

We then returned to the V2R, focusing on a mutant receptor truncated in its C-terminal cytoplasmic tail. This mutant V2R (HA-V2R-T) retains the ability to activate Gs but lacks phosphorylation sites which promote stable interaction with β-arrestins, causing the mutant receptor to internalize less efficiently after agonist-induced activation (Innamorati et al., 1998, 1997; Oakley et al., 1999). We confirmed this phenotype (**Fig 3C, 3D, 3E**) but observed, nevertheless, that HA-V2R-T was able to strongly stimulate internalization of Flag-AC9 (**Fig 3C, D, E** and **S3F**). This result, also verified by surface biotinylation (**Fig 3F, 3G**), supports the hypothesis that AC9 trafficking does not necessarily occur in complex with its upstream activating GPCR. Rather, it suggests that trafficking of AC9 and the activating GPCR are regulated separately.

### AC9 trafficking is not dependent on cytoplasmic cAMP or adenylyl cyclase activity

One possible mechanism of AC9 traffic control is by downstream cytoplasmic cAMP elevation stimulated by Gs. To test this, we applied the diterpene drug forskolin (FSK) to stimulate cytoplasmic cAMP production independently from the receptor or Gs. While AC9 is relatively insensitive to activation by FSK, other AC isoforms that are major contributors to cAMP production in HEK293 cells (such as AC3 and AC6) are sensitive, making FSK an effective stimulus of overall cAMP elevation (Baldwin et al., 2019; Soto-Velasquez et al., 2018). FSK did not cause detectable internalization of either HA-β2AR or Flag-AC9 assessed by imaging (**Fig S4A, S4C, S4D, S4l**) or surface biotinylation (**Fig S4E, S4F**). Further, as expected, Flag-AC1 remained in the plasma membrane (**Fig S4B, S4C, S4D, S4J**) irrespective of the presence of FSK (**Fig S4E, S4F**). Moreover, this was also the case after adding 3-isobutyl-1-methylxanthine (IBMX), a broad spectrum phosphodiesterase inhibitor known to additionally increase FSK-induced cAMP elevation in the cytoplasm (**Fig S4H**).

As an independent approach, and to consider the possibility that more localized cAMP production might be required, we asked if regulated trafficking of AC9 requires its own catalytic activity. To test this, we mutated a conserved aspartic acid residue that coordinates a catalytic magnesium in the active site, and which is essential for activity of AC6 (Gao et al., 2011; Tesmer et al., 1997). Mutating the equivalent residue in AC9 (Flag-AC9-D442A) blocked cAMP production (**Fig S1B**), but regulated trafficking of Flag-AC9-D442A was still observed (**Fig S4G**). Together, these results indicate that the ability of GPCR-Gs activation to regulate AC9 trafficking is not a consequence of global cytoplasmic cAMP elevation or of local cAMP production by AC9.

### Gs activation is sufficient to promote AC9 trafficking

We next investigated whether AC9 internalization is stimulated by Gs itself, and did so by introducing a point mutation into the alpha subunit (HA-Gs-Q227L) that increases Gs constitutive activity by reducing the rate of intrinsic GTP hydrolysis (Masters et al., 1989). Coexpressing Flag-AC9 with wild type HA-Gs resulted in localization of both proteins to the plasma membrane (**Fig 4A, 4C, 4D, S5E**). However, coexpression with activated HA-Gs-Q227L resulted in the localization of both proteins to internal punctae (**Fig 4A, 4C, 4D, S5E**). This effect was specific to AC9 because AC1 remained at the plasma membrane when coexpressed with either HA-Gs or HA-Gs-Q227L (**Fig 4B, 4C, 4D, S5F**). We verified by immunoisolation that HA-Gs-Q227L and Flag-AC9 both accumulate in EEA1-positive endosomes when coexpressed (**Fig S5A**), whereas Flag-AC1 did not accumulate in endosomes or cause Gs to accumulate there (**Fig S5B**). Independently supporting a discrete regulatory effect of Gs, application of cholera toxin (CTX) to activate the endogenous cellular complement of Gs resulted in receptor-independent accumulation of Flag-AC9, but not of AC1, in endosomes (**Fig S6**).

**Figure 4:**
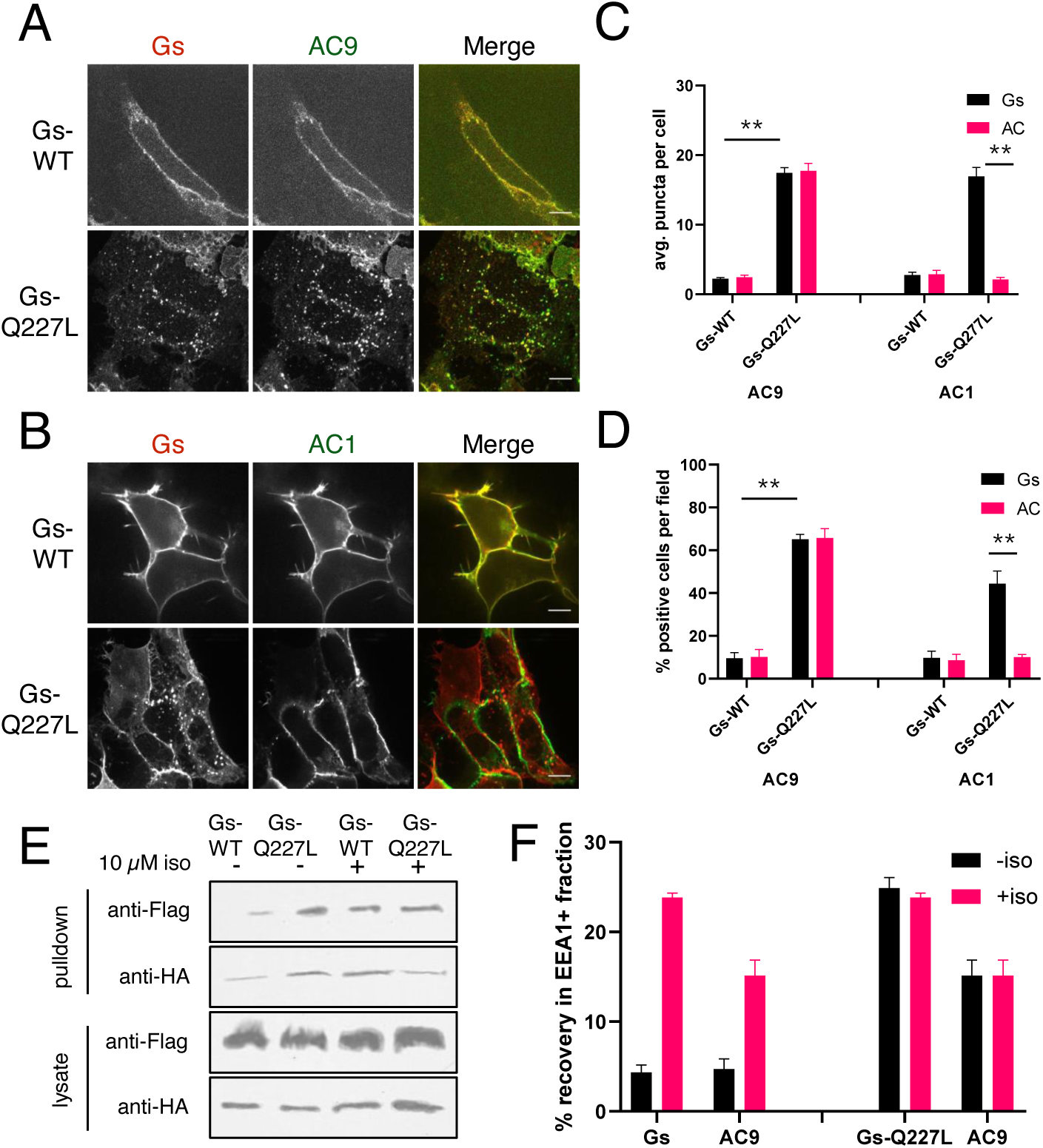
Gs activation is sufficient for AC9 to localize to early endosomes. **(A)** Representative confocal imaging of HEK293 cells cells coexpressing Flag-AC9, G_s_β, G_s_γ and either HA-G_s_α (HA-Gs) or HA-G_s_α-CA (HA-GsCA). **(B)** Representative confocal imaging of HEK293 cells coexpressing Flag-AC1, G_s_β, G_s_γ and either HA-Gsa (HA-Gs) or HA-G_s_α-CA (HA-GsCA). **(C)** Quantification of internal puncta that are Gs or AC1/9 positive, taken from wide field images (see: Supp. Fig 5C, 5D) [mean±SEM; n=3 experiments, 10 visual fields and 200+ cells per condition]. ** P < 0.01 by two-tailed t-test. **(D)** Quantification of cells with >10 internal puncta that are Gs or AC1/9 positive, taken from wide field images (see: Supp. Fig 5C, 5D) [mean±SEM; n=3 experiments, 10 visual fields and 200+ cells per condition]. ** P < 0.01 by two-tailed t-test. **(E)** Representative western blot of an EEA1+ fraction from HEK293 cells coexpressing Flag-AC9 and HA-Gs (lanes 1 and 3) or Flag-AC9 and HA-GsCA (lanes 2 and 4) and after treatment with 10 µM isoproterenol (lanes 1-2) or control (lanes 3-4) for 30 min. **(F)** Quantification of the fraction of Flag-AC9 and HA-Gs/HA-GsCA recovered in the EEA1+ fraction **(E)** relative to total cell lysate. [mean±SEM; n=3 experiments] * P < 0.05 ** P < 0.01 by two-tailed t-test.

We further confirmed HA-Gs and Flag-AC9 accumulation in EEA1-marked endosomes by immunoisolation (**Fig 4E, 4F**). Further, AC9 accumulation driven by activated Gs (expression of HA-Gs-Q227L) was not detectably increased by application of isoproterenol (**Fig 4E, 4F**). This suggests that Gs activation is fully sufficient to stimulate AC9 trafficking, without a requirement for additional effects of upstream receptor activation or downstream cAMP signaling.

### AC9 trafficking requires Gs but not β-arrestin

Since Gs activation is sufficient to stimulate the trafficking of AC9 to endosomes, we next asked if it is necessary for this regulated trafficking effect. To do so, we used previously described Gs-knockout (GsKO) cells that lack Gs due to CRISPR-mediated knockout of the Gs alpha subunit (Stallaert et al., 2017). Flag-β2AR and AC9-EGFP localized to the plasma membrane of GsKO as well as wild type HEK293 cells. However, AC9-GFP internalization was blocked in GsKO cells while Flag-β2AR was still internalized (**Fig 5A, 5C, 5D, S7A**). Moreover, AC9 trafficking was rescued by expression of recombinant HA-Gs (**Fig 5B, 5C, 5D, S7B**). These results indicate that Gs is necessary for regulated endocytic trafficking of AC9 but not of β2AR.

**Figure 5:**
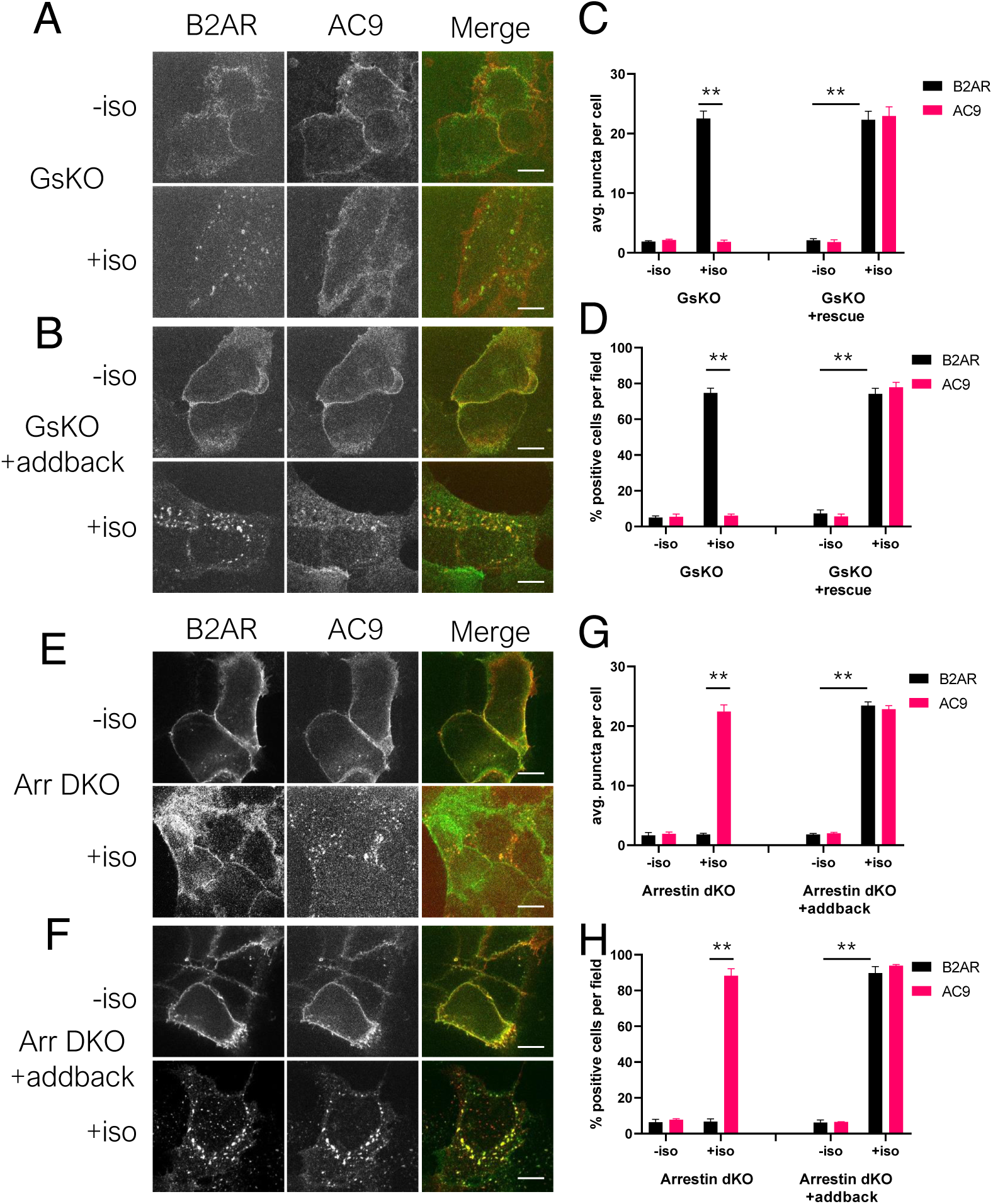
Gs activation is necessary for arrestin-independent endocytosis of AC9. **(A-B)** Representative confocal imaging of Gs knockout (GsKO) HEK293 cells coexpressing Flag-β2AR, AC9-EGFP, and either pcDNA3 **(A)** or wild-type HA-Gs rescue **(B)**. **(C-D)** Representative confocal imaging of Arrestin2/3 double knockout (Arr dKO) cells coexpressing Flag-β2AR, AC9-EGFP, and either pcDNA3 **(C)** or HA-Arrestin 3 rescue **(D)**. **(E)** Quantification of internal puncta that are β2AR or AC9 positive, taken from wide field images (see: Supp. Fig 7A, 7B) [mean±SEM; n=3 experiments, 10 visual fields and 200+ cells per condition]. ** P < 0.01 by two-tailed t-test. **(F)** Quantification of cells with >10 internal puncta that are β2AR or AC9 positive, taken from wide field images (see: Supp. Fig 7A, 7B) [mean±SEM; n=3 experiments, 10 visual fields and 200+ cells per condition]. ** P < 0.01 by two-tailed t-test. **(G)** Quantification of internal puncta that are β2AR or AC9 positive, taken from wide field images (see: Supp. Fig 7C, 7D) [mean±SEM; n=3 experiments, 10 visual fields and 200+ cells per condition]. ** P < 0.01 by two-tailed t-test. **(H)** Quantification of cells with >10 internal puncta that are Gs or AC1/9 positive, taken from wide field images (see: Supp. Fig 7C, 7D) [mean±SEM; n=3 experiments, 10 visual fields and 200+ cells per condition]. ** P < 0.01 by two-tailed t-test.

Because regulated entry of β2AR into the early endocytic pathway requires -βarrestins (Ferguson et al., 1996; Goodman et al., 1996), we asked if this is true also for AC9. To test this, we used HEK293 cells lacking both -βarrestin isoforms (Arrestins 2 and 3, or -βarrestin-1 and -βarrestin-2) generated using CRISPR (O’Hayre et al., 2017). Isoproterenol-stimulated internalization of HA-β2AR was blocked in these β-arrestin double-knockout (Arr DKO) cells, as expected, but AC9-EGFP was still accumulated in endosomes (**Fig 5E, 5G, 5H, S7C**). Moreover, expressing recombinant Arrestin 3 (β-arrestin-2) rescued the regulated trafficking of HA-β2AR without affecting AC9-EGFP (**Fig 5E, 5G, 5H, S7D**). These results indicate that regulated trafficking of AC9 in the shared early endocytic pathway is dependent on Gs but not β-arrestin, in contrast to regulated trafficking of the β2AR which is dependent on β-arrestin but not Gs.

### Functional evidence for AC9 signaling from endosomes

Both AC1 and AC9 are known physiological effectors of beta-adrenergic signaling (Sadana et al., 2009; Small et al., 2003; Tantisira, 2005) and both are endogenously expressed in HEK293 cells, despite neither being the primary contributor to global cAMP elevation produced by β2AR activation in this cell type. Nevertheless, siRNA-mediated knockdown analysis indicated that both AC1 and AC9 make a small but statistically significant contribution to the overall cAMP response elicited by endogenous β2ARs (**Fig S8A**). In contrast, AC1 but not AC9 knockdown reduced the FSK-induced cAMP response (**Fig S8B**), consistent with AC9 being resistant to stimulation by FSK (Baldwin et al., 2019) and verifying specificity of the knockdowns. Considering that AC9 selectively accumulates in endosomes relative to AC1, we next investigated the hypothesis that AC9 accordingly contributes to the endosome-initiated component of the β2AR-elicited cAMP response.

We tested this using a pharmacological approach, taking advantage of the ability of the membrane-impermeant β2AR antagonist CGP12177 (CGP) to access receptors only at the plasma membrane, while the membrane-permeant antagonist alprenolol accesses receptors both at the plasma membrane and endosomes (Staehelin et al., 1983). This strategy has been used successfully to isolate effects of endosome-localized receptor activation by comparative blockade (Irannejad et al., 2013; Thomsen et al., 2016). We validated this strategy in our hands using a conformational biosensor, Nb80-EGFP, to detect the reversible activation of β2ARs (Irannejad et al., 2013). Isoproterenol application produced recruitment of Nb80-GFP by β2ARs both at the plasma membrane and endosomes, and subsequent addition of excess alprenolol rapidly reversed Nb80-GFP recruitment at both locations (**Fig 6A**, Supplemental Video 2). CGP, however, reversed recruitment at the plasma membrane but not endosomes (**Fig 6B**, Supplemental Video 3).

**Figure 6:**
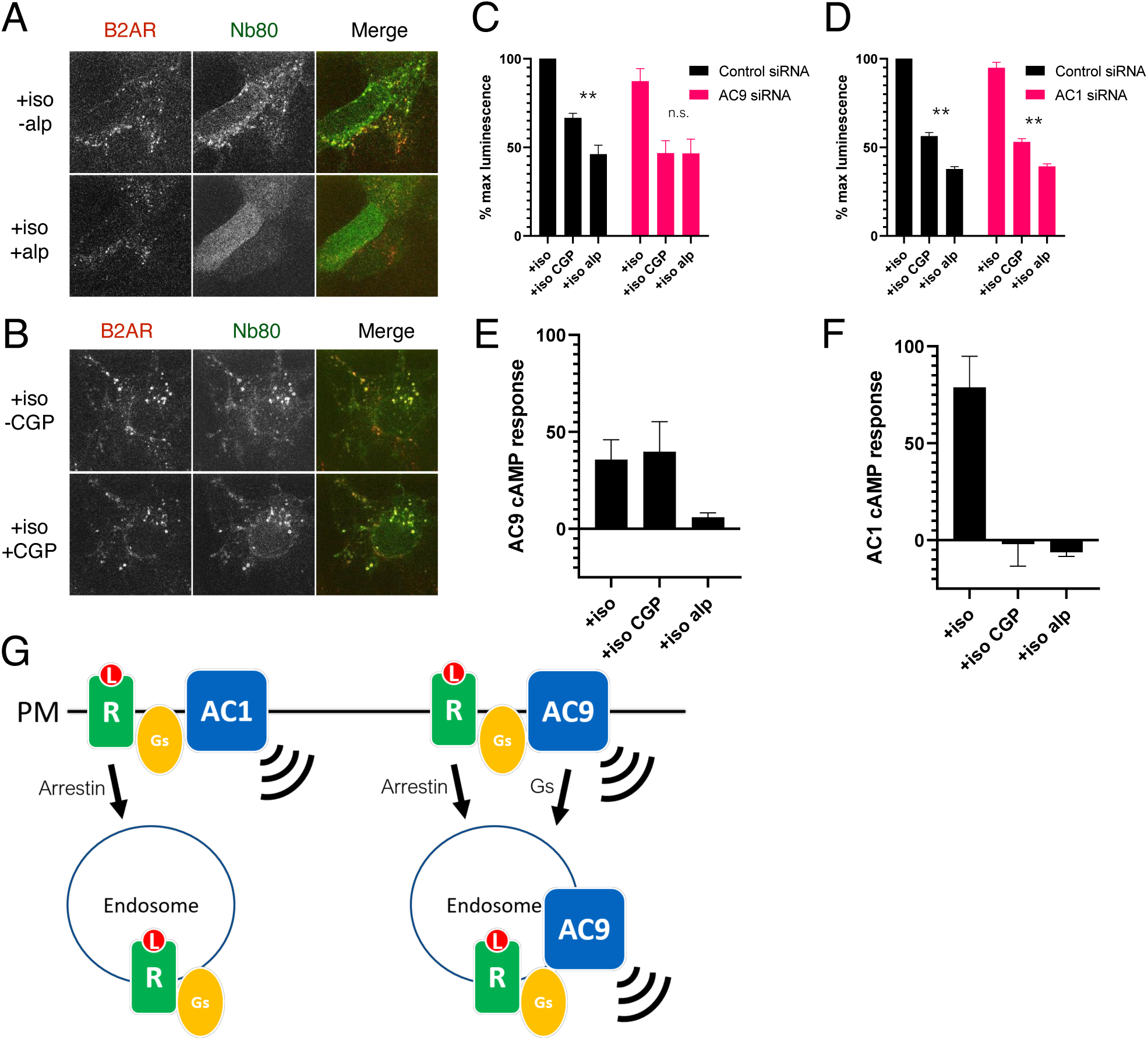
AC9 is a significant contributor to the β2AR-mediated cAMP response from endosomes. **(A)** Recruitment of conformational biosensors to β2AR-containing endosomes is reversed by application of the membrane permeable antagonist alprenolol for 20 min. Scale Bar is 8 µm. See Supplemental Video 2 for full image series. **(B)** Recruitment of conformational biosensors to β2AR-containing endosomes is unaffected by application of the membrane impermeable antagonist CGP12177 for 20 min. Scale Bar is 8 µm. See Supplemental Video 3 for full image series. **(C)** Quantification of the maximum cAMP response in control and in AC9 siRNA knockdown HEK293 cells pretreated with 100nM isoproterenol and exposed to supersaturating conditions of membrane permeable antagonist (10µM alprenolol) or membrane impermeable antagonist (10µM CGP12177). [mean±SEM; n=4 experiments] **(D)** Quantification of the maximum cAMP response in control in AC1 siRNA knockdown HEK293 cells pretreated with 100nM isoproterenol and exposed to supersaturating conditions of membrane permeable antagonist (10µM alprenolol) or membrane impermeable antagonist (10µM CGP12177). [mean±SEM; n=4 experiments] **(E)** Quantification of the maximum cAMP response in AC3/6KO HEK293 cells due to AC9 overexpression as the delta between the Flag-AC9 and pcDNA3 conditions. Cells were pretreated with 100nM isoproterenol and exposed to supersaturating conditions of membrane permeable antagonist (10µM alprenolol) or membrane impermeable antagonist (10µM CGP12177). [mean±SEM; n=4 experiments] **(F)** Quantification of the maximum cAMP response in AC3/6KO HEK293 cells due to AC1 overexpression as the delta between the Flag-AC1 and pcDNA3 conditions. Cells were pretreated with 100nM isoproterenol and exposed to supersaturating conditions of membrane permeable antagonist (10µM alprenolol) or membrane impermeable antagonist (10µM CGP12177). [mean±SEM; n=4 experiments] **(G)** Model: Ligand binding causes initial signaling event at the PM followed by arrestin-dependent endocytosis of β2AR. AC1 is restricted to the PM but AC9 is dynamically redistributed by a distinct Gs regulated process, and contributes to the β2AR-mediated cAMP response from the endosome.

We next applied this strategy to probe the contribution of β2AR activation in endosomes to the overall cellular cytoplasmic cAMP response. Both CGP and alprenolol markedly reduced cytoplasmic cAMP elevation produced by isoproterenol and measured using a real-time cAMP biosensor (Irannejad et al., 2013). This verifies that a large fraction of the overall cAMP elevation elicited by endogenous β2AR activation in HEK293 cells is indeed initiated from the plasma membrane. However, in addition, we consistently observed a more pronounced inhibition of the cellular cAMP response following application of excess alprenolol compared to CGP (**Fig 6C**, left set of bars; see also **Fig S8G** and Supplemental Video 4). We interpret this difference as a readout of the component of cAMP production initiated by β2AR activation in endosomes. Remarkably, this CGP-resistant ‘signal gap’ was lost after AC9 knockdown (**Fig 6C**) but it remained in cells depleted of AC1 (**Fig 6D**). Nevertheless, both knockdowns produced a similar reduction of the overall isoproterenol-induced cytoplasmic cAMP elevation measured in the absence of either antagonist (**Fig S8B**).

As another test of this hypothesis, and to investigate selectivity under conditions of recombinant AC overexpression (which were necessarily used for the trafficking studies), we asked if a specific endosomal signaling effect is evident also using tagged AC isoforms. To do so, we used HEK293 cells depleted of both AC3 and AC6 (AC3/6 DKO) that were shown previously to provide a reduced background useful for assessing effects of recombinant AC expression on cellular cAMP (Soto-Velasquez et al., 2018). The increase of isoproterenol-induced cAMP accumulation produced by overexpression of Flag-AC9, defined as the ‘AC9 response’, was blocked by alprenolol but not CGP (**Fig 6E**). However, the AC1 response, assessed in the same way using Flag-AC1, was blocked both by alprenolol and CGP (**Fig 6F**). Together these results verify that AC9 contributes specifically to the cellular cAMP response initiated by β2AR activation in endosomes, and suggest that this is true for recombinant as well as endogenous AC9.

## Discussion

The endocytic network is a dynamically regulated system critical for homeostatic integrity of the cell. From the point of view of GPCR-G protein signaling, this network was believed for many years to be silent, and to function only in signal termination and longer-term modulation of surface receptor number. Such homeostatic effects indeed occur, but an accumulating body of evidence supports an expanded view in which internalized GPCRs reacquire the ability to activate G proteins after endocytosis and initiate a second wave of signaling from endomembrane sites (Irannejad et al., 2015; Lohse and Calebiro, 2013; Vilardaga et al., 2014). Endosomal signaling depends on the presence of a G protein-regulated effector, but whether or how effectors localize to relevant internal membrane locations has remained a relatively unexplored frontier.

We approached this problem by focusing on ACs as important effectors of signaling initiated by GPCR - Gs activation. We demonstrate dynamic and regulated trafficking of AC9 to early endosomes that are known to accumulate a wide variety of GPCRs, and which have been explicitly shown to be a site of Gs activation by several GPCRs. AC9 is a physiologically and clinically relevant effector of β2AR-Gs signaling in particular (Small et al., 2003; Sunahara, 1996; Tantisira, 2005), and it is endogenously expressed in the HEK293 model system used in the present study. We also show that AC9, while contributing only a minor fraction to the overall cellular cAMP response elicited by β2AR activation in this system, is necessary for producing a specific endosome-initiated component of the endogenous cAMP response. We show, further, that AC9 is sufficient to increase cAMP production from endosomes when expressed as a recombinant protein. Moreover, we show that AC trafficking is isoform-specific because AC1 does not detectably accumulate in endosomes, nor does AC1 contribute detectably to the endosome-initiated component of cellular cAMP signaling.

An important future goal is to identify structural and biochemical determinants of isoform-specific AC trafficking. We note that various isoform-specific protein interactions which impact other aspects of AC function are already known, with AC9 being a particularly well-studied example (Baldwin et al., 2019). Another important question for future investigation is whether regulated intracellular trafficking is unique to AC9 or is more widespread. We anticipate the latter because a distantly related AC isoform has been previously localized to a multivesicular intracellular compartment in *D. discoideum* (Kriebel et al., 2008). However, in this case, trafficking appears to occur through the biosynthetic pathway and it is not known if the AC-containing compartment also contains a relevant GPCR or G protein. We also note that AC2 and AC3 contribute to cAMP signaling by Gs-coupled polypeptide hormone receptors after endocytosis in mammalian cells (Jean-Alphonse et al., 2017; Kriebel et al., 2008), but the trafficking properties of these AC isoforms remain to be delineated. Moreover, a distinct AC isoform that lacks any transmembrane domains (‘soluble’ AC or AC10) contributes to the cellular cAMP response elicited by another Gs-coupled GPCR from endosomes (Inda et al., 2016). Accordingly, we anticipate that there likely exist multiple mechanisms for spatiotemporal control of cAMP signaling, from endosomes as well as from other endomembrane compartments (Irannejad et al., 2017).

One possible mechanism for trafficking AC to GPCR-containing endosomes is by physical association with the receptor or receptor-G protein complex, and there is evidence for such complex formation involving AC5 (Navarro et al., 2018). Our results indicate that AC9, despite traversing a similar dynamin-dependent membrane pathway as the β2AR, traffics independently. First, activation of Gs is sufficient to promote AC9 but not β2AR accumulation in endosomes. Second, AC9 trafficking requires Gs but not β-arrestins, whereas the converse is true for trafficking of the β2AR. These results indicate that AC9 trafficking in the endocytic network, while mediated by the same core pathway as that mediating GPCR internalization, and despite delivering both proteins to the same endosome membranes, differs fundamentally in its mechanism of regulation. Accordingly, AC trafficking is likely subject to different modulatory input(s) relative to the trafficking of GPCRs. This is consistent with the particular environmental sensitivity of AC9 relative to β2AR trafficking that motivated our initial studies. However, additional study will be required to define the mechanistic basis for differential control of AC9, and to delineate physiological input(s) regulating AC trafficking more broadly. The functional significance of distinct AC traffic control also remains to be further investigated, but we note that there is already significant evidence that cAMP production internally in the cytoplasm can produce different downstream effects relative to production from the plasma membrane (O’Banion et al., 2019; Tsvetanova and von Zastrow, 2014).

In closing, to our knowledge the present study is the first to delineate the dynamic endocytic trafficking of a functionally relevant AC isoform, and to identify a role of Gs in regulating the trafficking of an AC separately from its catalytic activity. The finding that AC9 trafficking is isoform-specific, and regulated separately from its activating GPCR, reveals a new layer of specificity and control in the cAMP cascade.

## Acknowledgements

We thank B. Lobingier, Grace Peng, Lea Ripoll, M. Stoeber and other current and past members of the von Zastrow laboratory for useful advice and critical discussions. We thank J. Taunton, D. Mullins and R. Sunahara for valuable discussion and suggestions. We thank Monica Soto-Velasquez and V. Watts for valuable discussion and the generous gift of AC3/6 knockout cells. We thank the E. Roth group for their generous support and lab equipment, and Kyra Kurtz for assistance. This work was supported by grants from the US National Institutes of Health (DA010711 and DA012864 to M.v.Z.; GM60419 to C.W.D; HL124049 to A.A.S; CA209891 to J.S.G.). A.L. is an ARCS scholar.

## Author contributions

A.L. conceived and designed experiments, performed all experiments, analyzed data, and wrote the paper. R.I. conceived and designed experiments. T.A.B. developed constructs and performed experiments. A.A.S. prepared primary cell culture human airway smooth muscle cells. C.W.D. provided necessary reagents and wrote the paper. A. I. provided necessary reagent and wrote the paper. M.v.Z. conceived and designed experiments, analyzed data, and wrote the paper.

## Declaration of interests

The authors declare no competing interests

## Figure legends

**Supplementary Figure 1.**
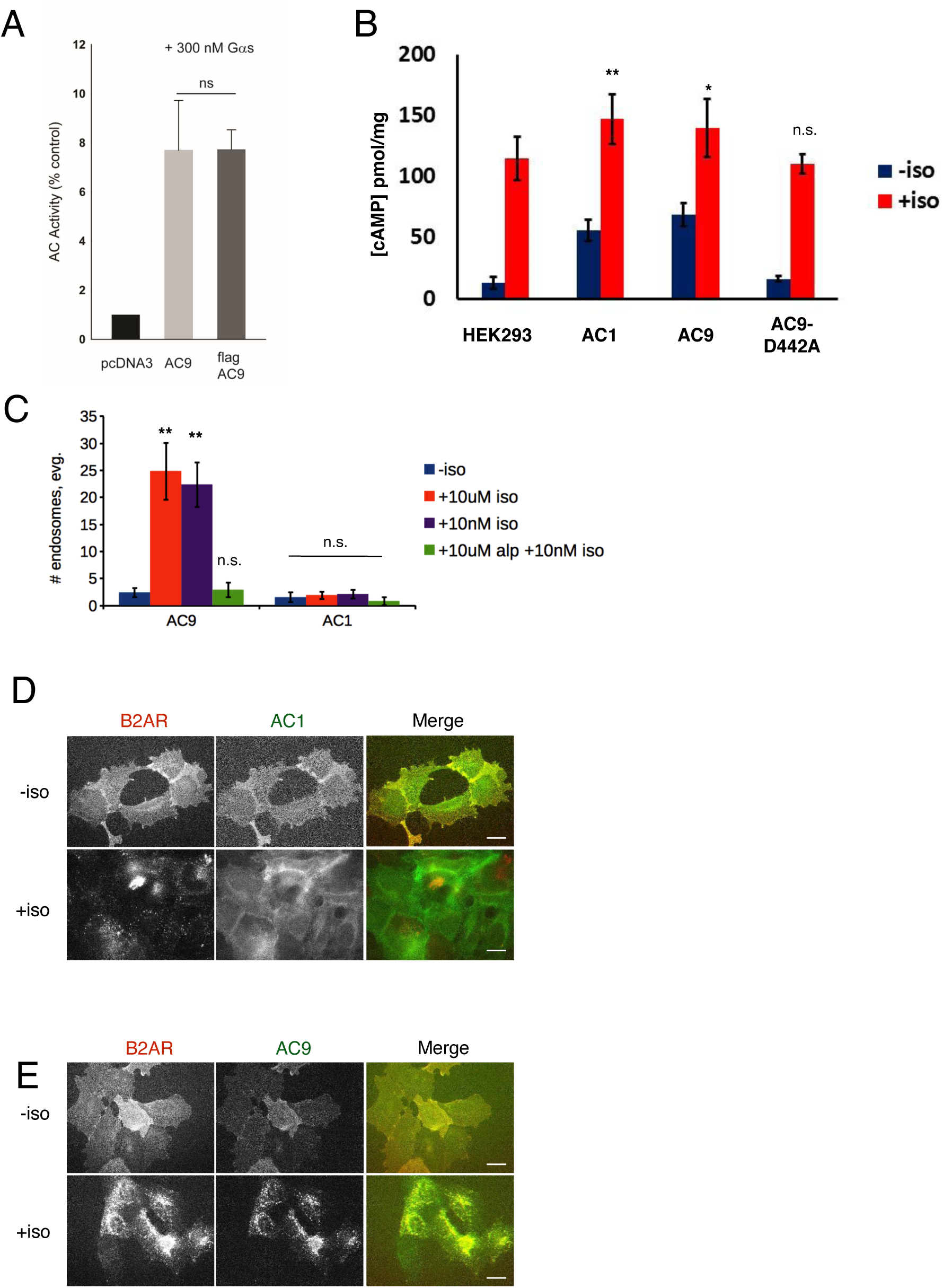
**(A)** Quantification of the in vitro cAMP response by pcDNA3 control, untagged AC9, and Flag-tagged AC9 to 300 nM Gs (alpha subunit). **(B)** Quantification of the cAMP response to 10µM isoproterenol in control HEK293 cells and cells overexpressing Flag-AC1, Flag-AC9, or Flag-AC9-D442A by ELISA assay. [mean±SEM; n=3 experiments] **(C)** Quantification of the number of endosomes in cells from **(Fig 1E-1H)** [mean±SEM; n=3 experiments, 25 cells per condition]. **(D)** Wide field images of HEK293 cells coexpressing HA-β2AR and Flag-AC9, after treatment with 10 µm isoproterenol or control for 30 min. Scale bar is 16 µm. **(E)** Wide field images of HEK293 cells coexpressing HA-β2AR and Flag-AC1, after treatment with 10 µm isoproterenol or control for 30 min. Scale bar is 16 µm.

**Supplementary Figure 2.**
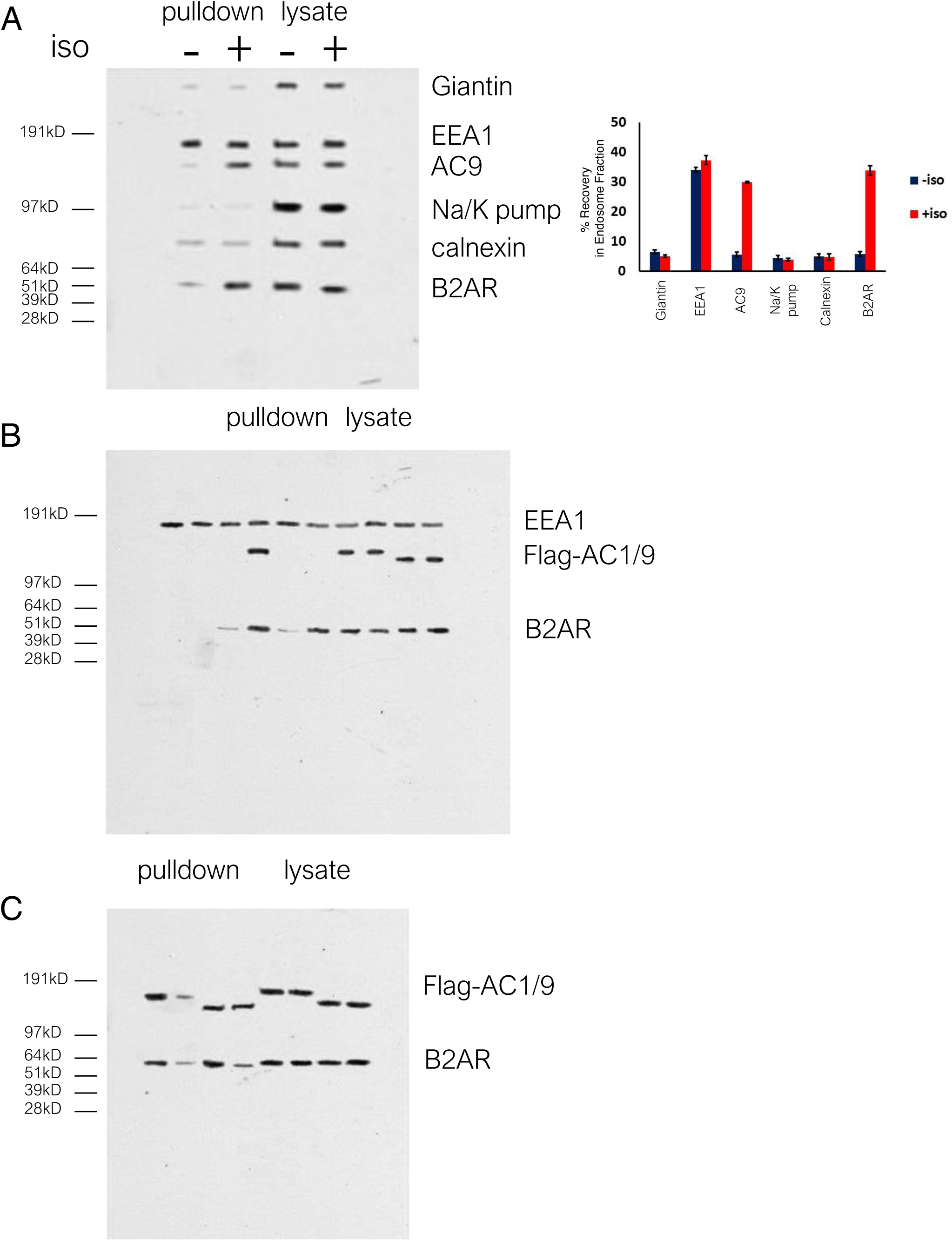
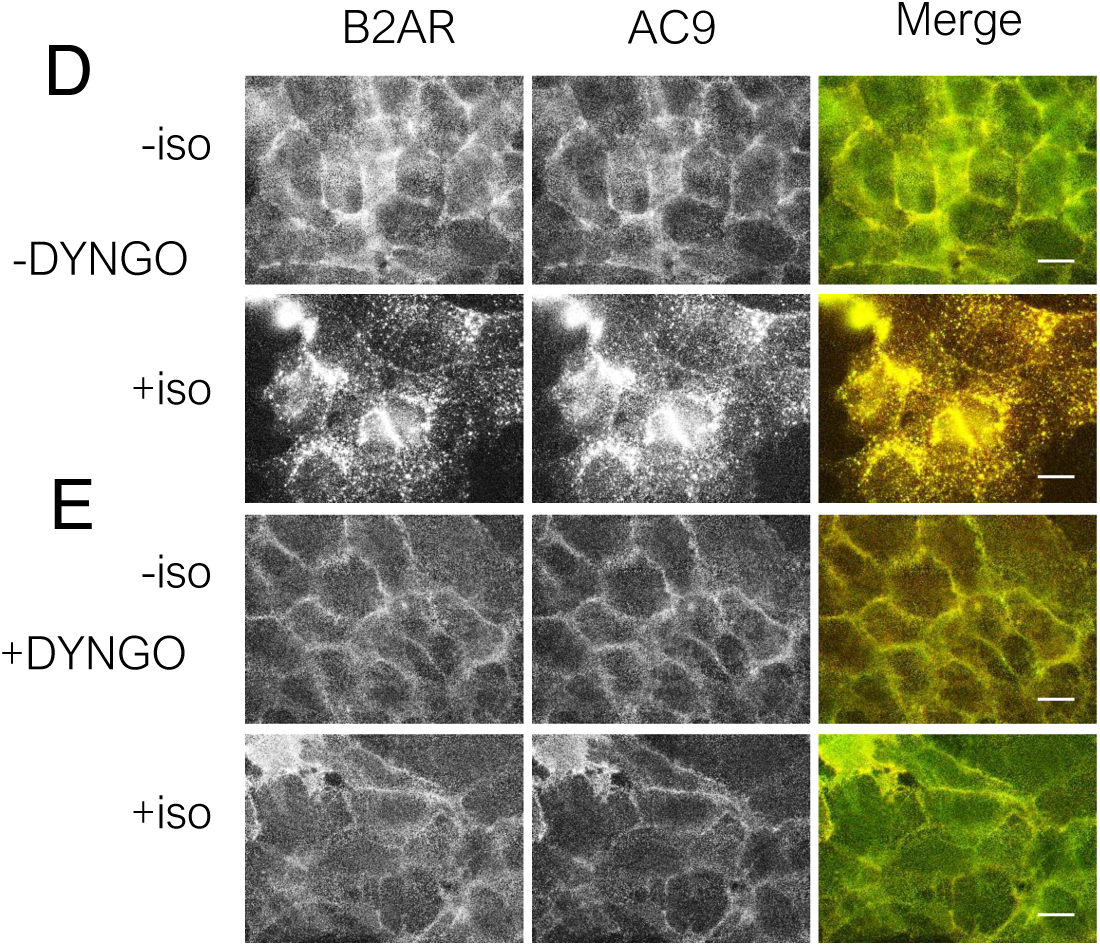
**(A)** Full western blot of the endosome fraction vs total cell lysate, probed for cellular markers Giantin (Golgi), Na/K pump (plasma membrane), calnexin (endoplasmic reticulum). Quantification of 3 such experiments. **(B)** uncut western blot from Figure 2C. **(C)** uncut western blot from Figure 2E. **(D)** Wide field images of HEK293 cells coexpressing HA-β2AR and Flag-AC9, pretreated with DMSO control for 15 min, after treatment with 10 µm isoproterenol or control for 30 min. Scale bar is 16 µm. **(E)** Wide field images of HEK293 cells coexpressing HA-β2AR and Flag-AC1, pretreated with 30uM DYNGO-4a, after treatment with 10 µm isoproterenol or control for 30 min. Scale bar is 16 µm.

**Supplementary Figure 3.**
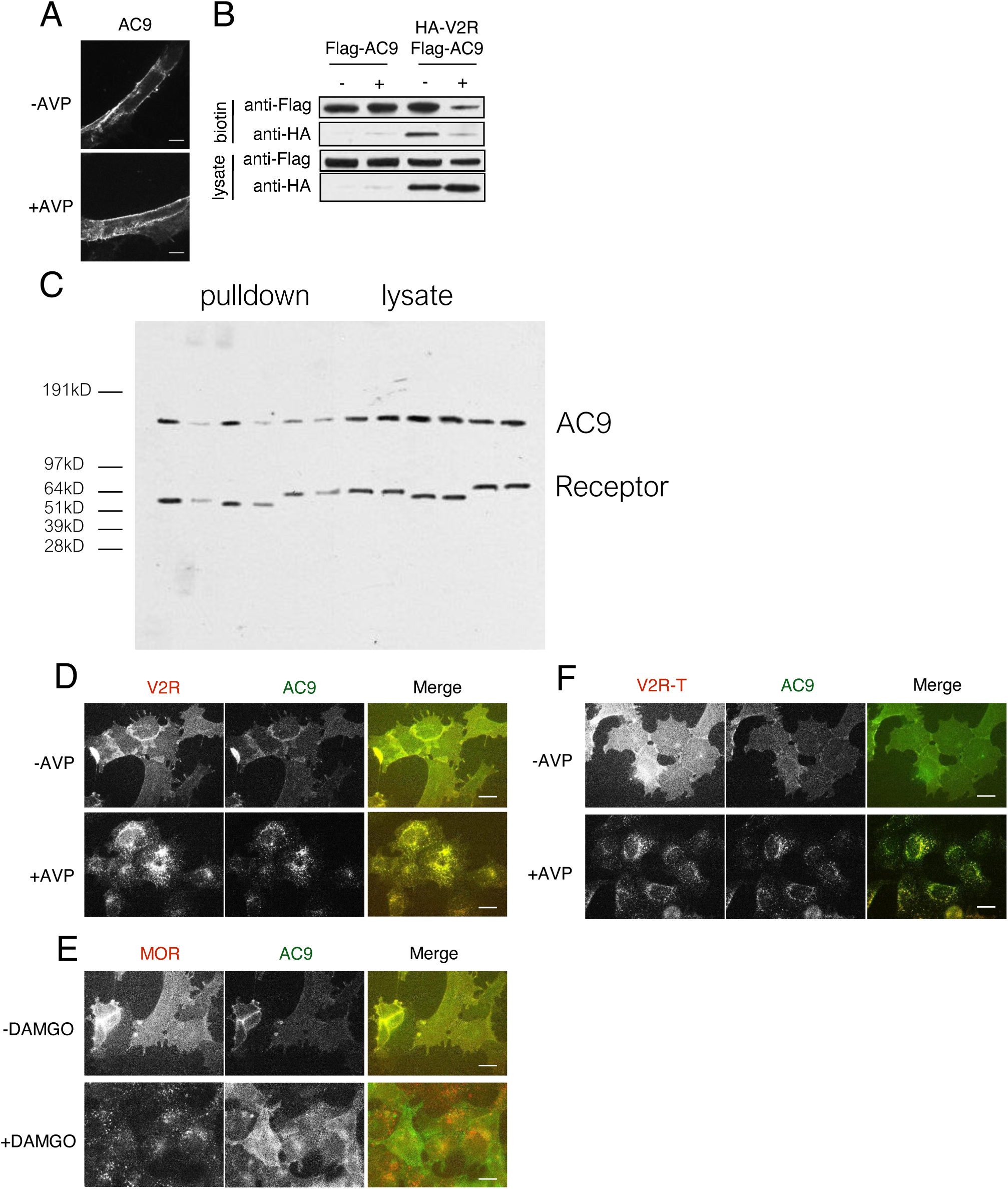
**(A)** Representative confocal imaging of HEK293 cells expressing Flag-AC9, after treatment with 10 µM AVP or control for 30 min. Scale Bar is 8 µm. **(B)** Representative western blot of the surface biotinylated fraction from HEK293 cells expressing Flag-AC9 (lanes 1-2) or coexpressing HA-V2R and Flag-AC9 (lanes 3-4) [mean±SEM; n=3 experiments]. **(C)** Uncut blot from Figure 3F. **(D)** Wide field images of HEK293 cells coexpressing HA-V2R and Flag-AC9, after treatment with 10 µm arginine vasopressin (AVP) or control for 30 min. Scale bar is 16 µm. **(E)** Wide field images of HEK293 cells coexpressing HA-MOR and Flag-AC9, after treatment with 10 µm [D-Ala^2^, N-MePhe^4^, Gly-ol]-enkephalin (DAMGO) or control for 30 min. Scale bar is 16 µm. **(F)** Wide field images of HEK293 cells coexpressing HA-V2R-trunc and Flag-AC9, after treatment with 10 µm arginine vasopressin (AVP) or control for 30 min. Scale bar is 16 µm.

**Supplementary Figure 4:**
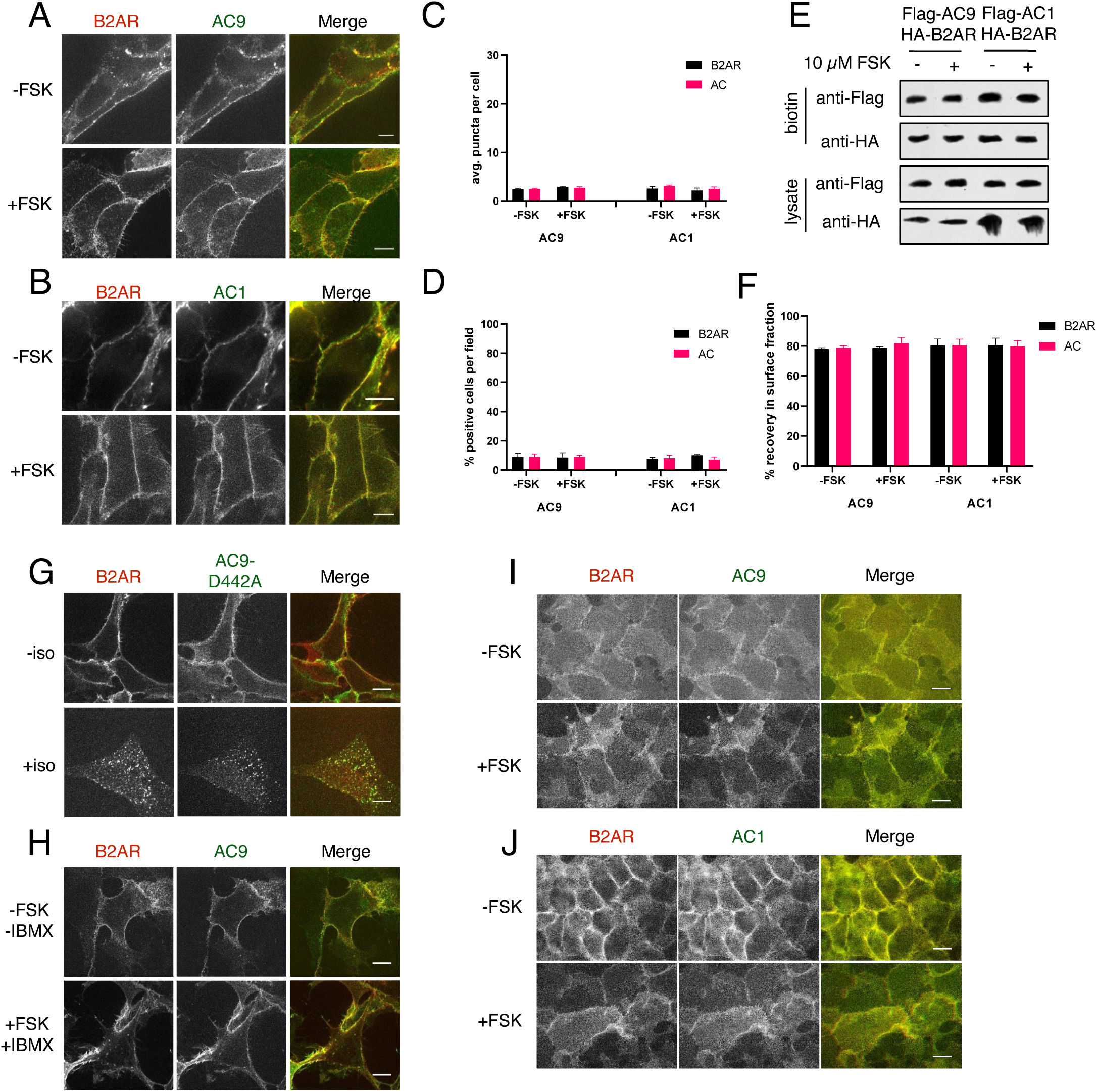
Forskolin-promoted cAMP accumulation is insufficient to drive AC9 internalization. **(A)** Representative confocal imaging of HEK293 cells coexpressing HA-β2AR and Flag-AC9 were treated with 10 µM FSK or control for 30 min. Scale Bar is 5 µm. **(B)** Representative confocal imaging of HEK293 cells coexpressing HA-β2AR and Flag-AC1 were treated with 10 µM FSK or control for 30 min. Scale Bar is 5 µm. **(C)** Quantification of internal puncta that are β2AR or AC1/9 positive, taken from wide field images (see: Supp. Fig 4I, 4J) [mean±SEM; n=3 experiments, 10 visual fields and 200+ cells per condition]. ** P < 0.01 by two-tailed t-test. **(D)** Quantification of cells with >10 internal puncta that are β2AR or AC1/9 positive, taken from wide field images (see: Supp. Fig 4I, 4J) [mean±SEM; n=3 experiments, 10 visual fields and 200+ cells per condition]. ** P < 0.01 by two-tailed t-test. **(E)** Representative western blot of the surface biotinylated fraction of cells from **(A**,**C)**. **(F)** Quantification of the percent loss from the surface biotinylated fraction relative to the total cell lysate as seen in **(E)** [mean±SEM; n=5 experiments]. **(G)** Representative confocal imaging of HEK293 cells coexpressing HA-β2AR and the catalytic inactive mutant Flag-AC9-D442A were treated with 10 µM isoproterenol or control for 30 min. Scale Bar is 5 µm. **(H)** Representative confocal imaging of HEK293 cells coexpressing HA-β2AR and Flag-AC9 were treated with 10 µM FSK and 10 µM IBMX or control for 30 min. Scale Bar is 5 µm. **(l)** Representative wide-field imaging of HEK293 cells coexpressing HA-β2AR and Flag-AC9 which were treated with 10 µM FSK or control for 30 min. Scale Bar is 16 µm. **(J)** Representative wide-field imaging of HEK293 cells coexpressing HA-β2AR and Flag-AC1 which were treated with 10 µM FSK or control for 30 min. Scale Bar is 16 µm.

**Supplementary Figure 5.**
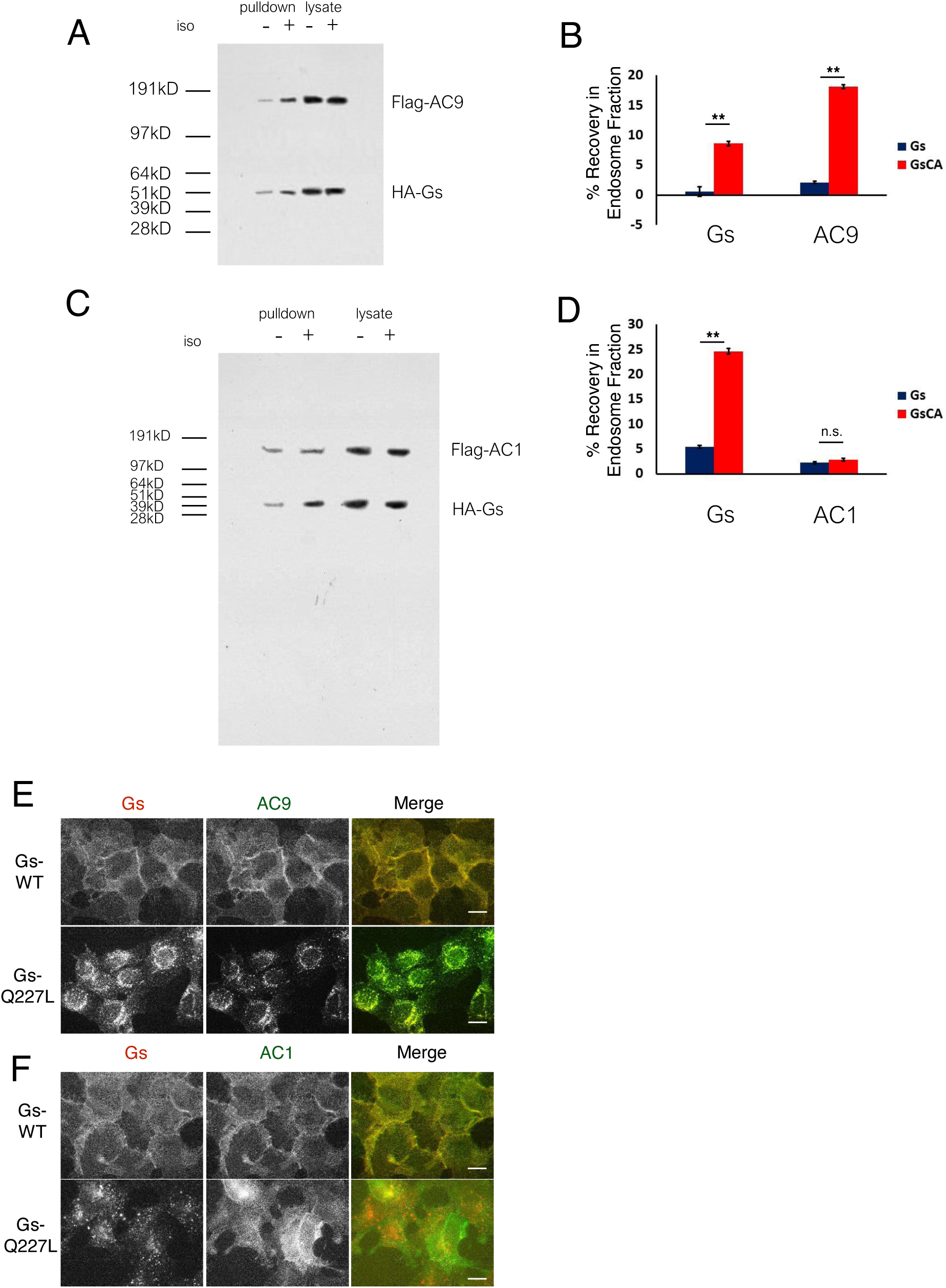
**(A)** Representative western blot of an EEA1 positive endosome fraction from HEK293 cells coexpressing Flag-AC9, G_s_β, G_s_γ and HA-Gs (lane 1) or HA-GsCA (lane 2). **(B)** Quantification of endosome enrichment of Flag-AC9 and HA-Gs or HA-GsCA as seen in **(A)** by normalizing to the EEA1 signal. [mean±SEM; n=5 experiments] ** P < 0.01 by two-tailed t-test. **(C)** Representative western blot of an EEA1 positive endosome fraction from HEK293 cells coexpressing Flag-AC1, G_s_β, G_s_γ and HA-Gs (lane 1) or HA-GsCA (lane 2). **(D)** Quantification of endosome enrichment of Flag-AC1 and HA-Gs or HA-GsCA as seen in **(B)** by normalizing to the EEA1 signal. [mean±SEM; n=5 experiments] ** P < 0.01 by two-tailed t-test. **(E)** Representative wide-field imaging of HEK293 cells coexpressing HA-Gs (wild type) or HA-Gs-Q227L and Flag-AC9. Scale Bar is 16 µm. **(F)** Representative wide-field imaging of HEK293 cells coexpressing HA-Gs (wild type) or HA-Gs-Q227L and Flag-AC1. Scale Bar is 16 µm.

**Supplementary Figure 6.**
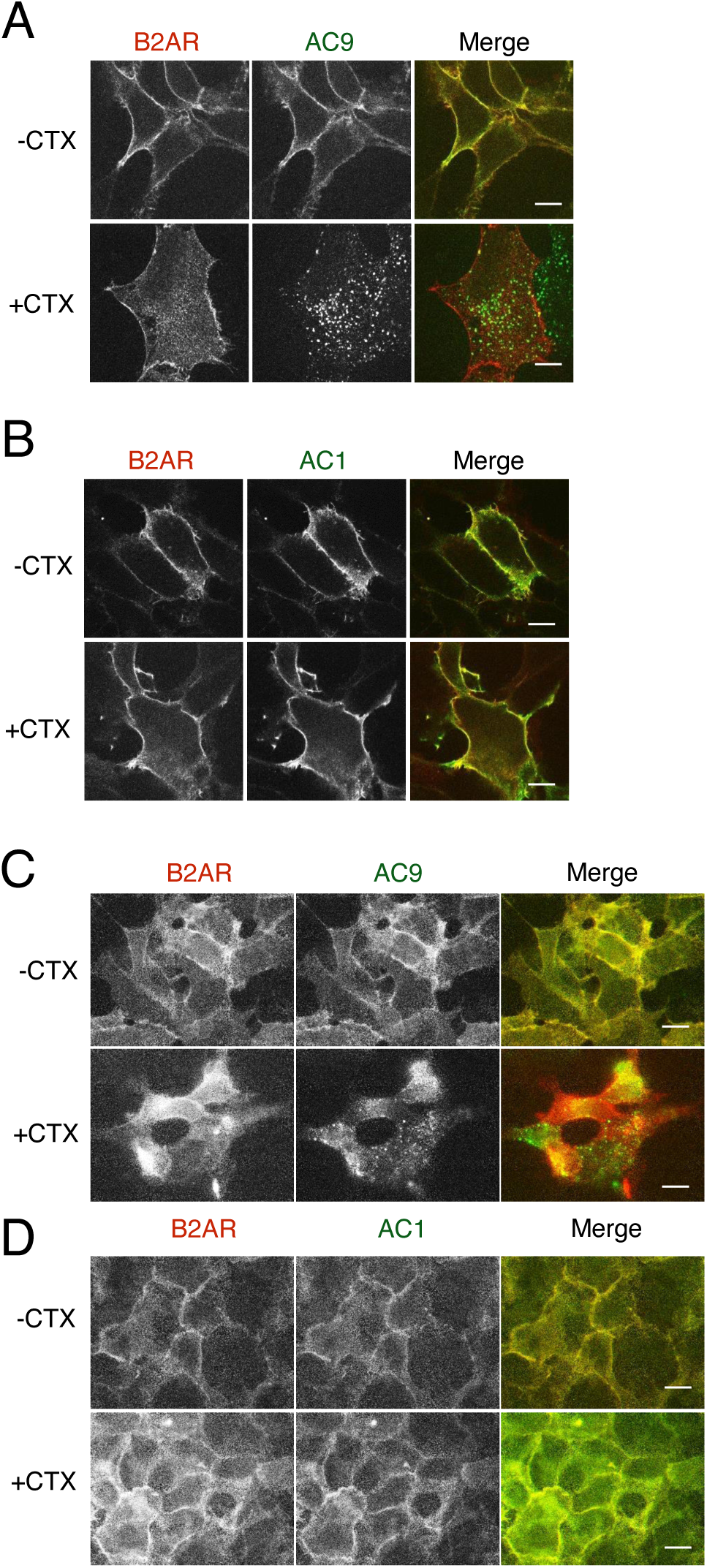
**(A)** Representative confocal imaging of HEK293 cells coexpressing HA-β2AR and Flag-AC9 were treated with 100 nM cholera toxin overnight. Scale Bar is 8 µm. **(B)** Representative confocal imaging of HEK293 cells coexpressing HA-β2AR and Flag-AC1 were treated with 100 nM cholera toxin overnight. **(C)** Representative wide-field imaging of HEK293 cells coexpressing HA-β2AR and Flag-AC9 were treated with 100 nM cholera toxin overnight. Scale Bar is 8 µm. **(D)** Representative wide-field imaging of HEK293 cells coexpressing HA-β2AR and Flag-AC1 were treated with 100 nM cholera toxin overnight.

**Supplementary Figure 7.**
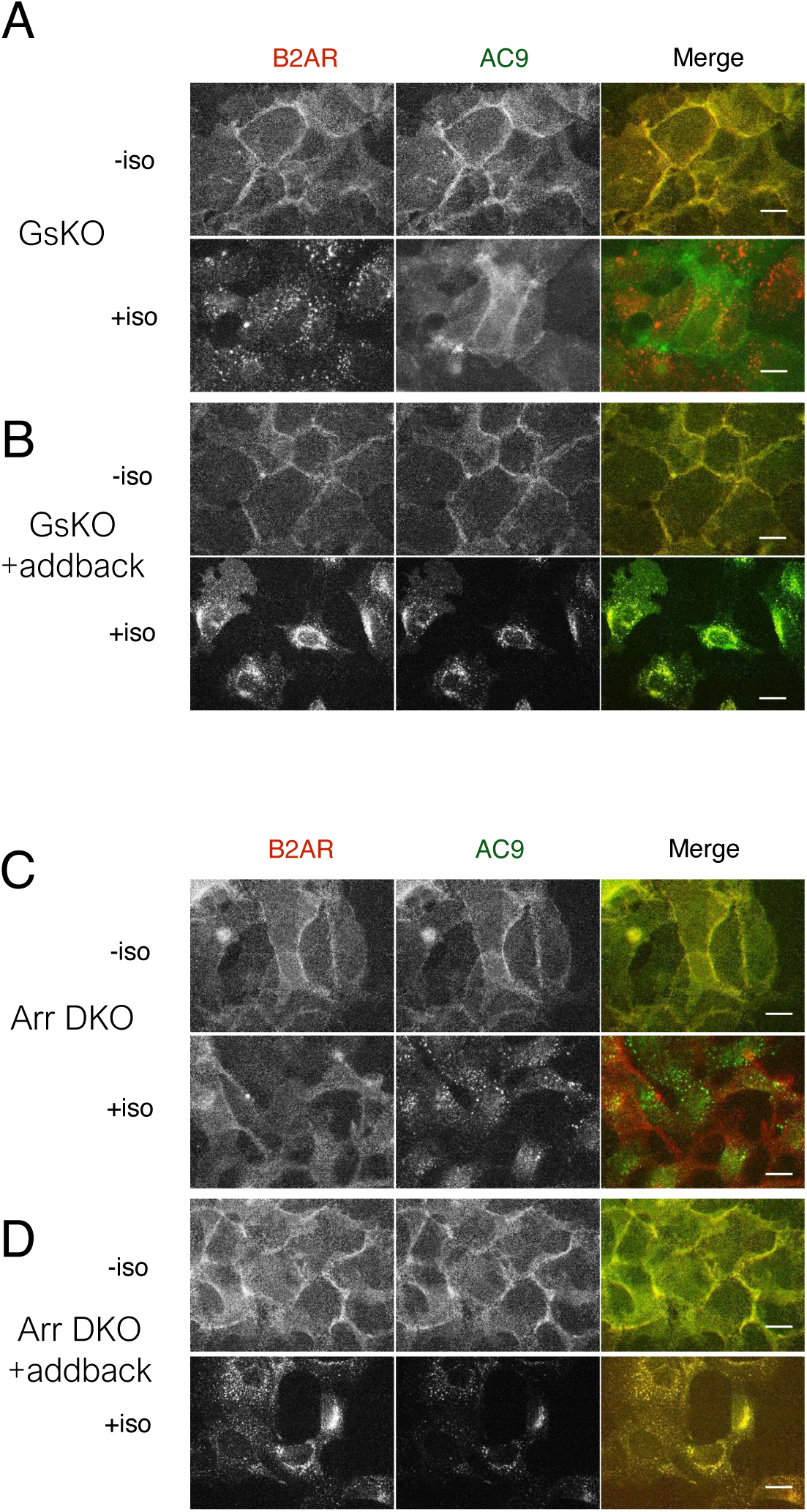
**(A)** Representative wide-field imaging of Gs-knockout HEK293 cells coexpressing HA-β2AR and Flag-AC9 were treated with 10 µM isoproterenol for 30 min. Scale Bar is 16 µm. **(B)** Representative wide-field imaging of Gs-knockout HEK293 cells coexpressing Gs (wild type) HA-β2AR and Flag-AC9 were treated with 10 µM isoproterenol for 30 min. Scale Bar is 16 µm. **(C)** Representative wide-field imaging of Arrestin 2/3 double knockout HEK293 cells coexpressing HA-β2AR and Flag-AC9 were treated with 10 µM isoproterenol for 30 min. Scale Bar is 16 µm. **(D)** Representative wide-field imaging of Arrestin 2/3 double knockout HEK293 cells coexpressing Arrestin 2, HA-β2AR and Flag-AC9 were treated with 10 µM isoproterenol for 30 min. Scale Bar is 16 µm.

**Supplementary Figure 8.**
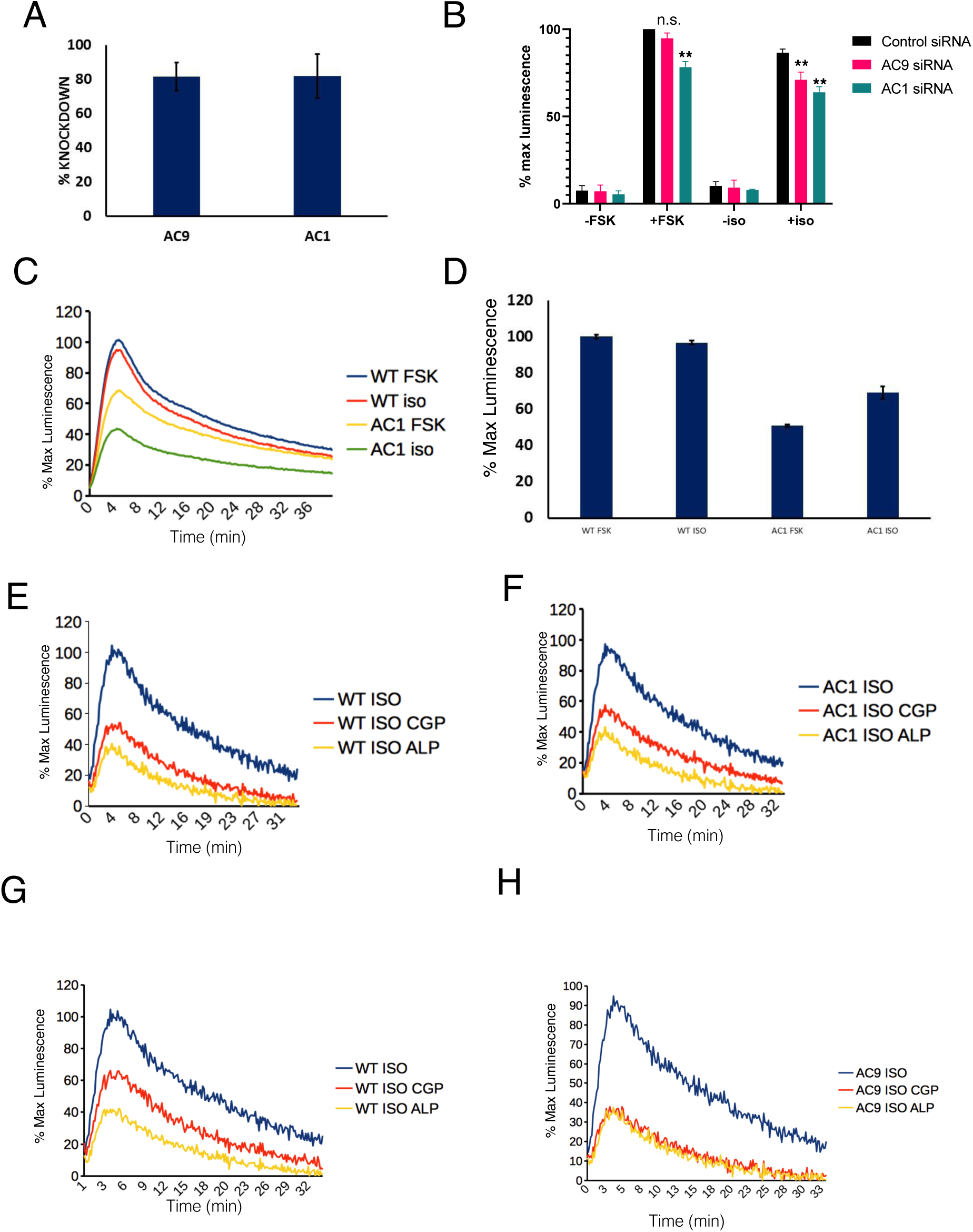
**(A)** Percent knockdown of AC9 and AC1 in HEK293 cells as determined by qPCR. [mean±SEM; n=3 experiments] **(B)** Effect of siRNA knockdown of AC9 and AC1 expression in HEK293 cells on the cAMP response to 10 µM FSK and 10 µM isoproterenol stimulation [mean±SEM; n=4 experiments] * P < 0.05 by two-tailed t-test. **(C)** Representative cAMP response to 10 µM FSK or 10 µM isoproterenol under conditions of siRNA knockdown of AC1 expression or control (ASD). **(D)** Quantification of the maximum cAMP response as seen in **(C)** [mean±SEM; n=3 experiments]. **(E)** Representative normalized β2AR-mediated cAMP response in control HEK293 cells pretreated with 100nM isoproterenol and exposed to supersaturating antagonist conditions (10µM CGP12177, 10µM alprenolol). **(F)** Representative normalized β2AR-mediated cAMP response in AC1 knockdown HEK293 cells pretreated with 100nM isoproterenol and exposed to supersaturating antagonist conditions (10µM CGP12177, 10µM alprenolol). **(G)** Representative normalized β2AR-mediated cAMP response in control HEK293 cells pretreated with 100nM isoproterenol and exposed to supersaturating antagonist conditions (10µM CGP12177, 10µM alprenolol). **(H)** Representative normalized β2AR-mediated cAMP response in AC9 knockdown HEK293 cells pretreated with 100nM isoproterenol and exposed to supersaturating antagonist conditions (10µM CGP12177, 10µM alprenolol).

**Supplementary Video 1.**
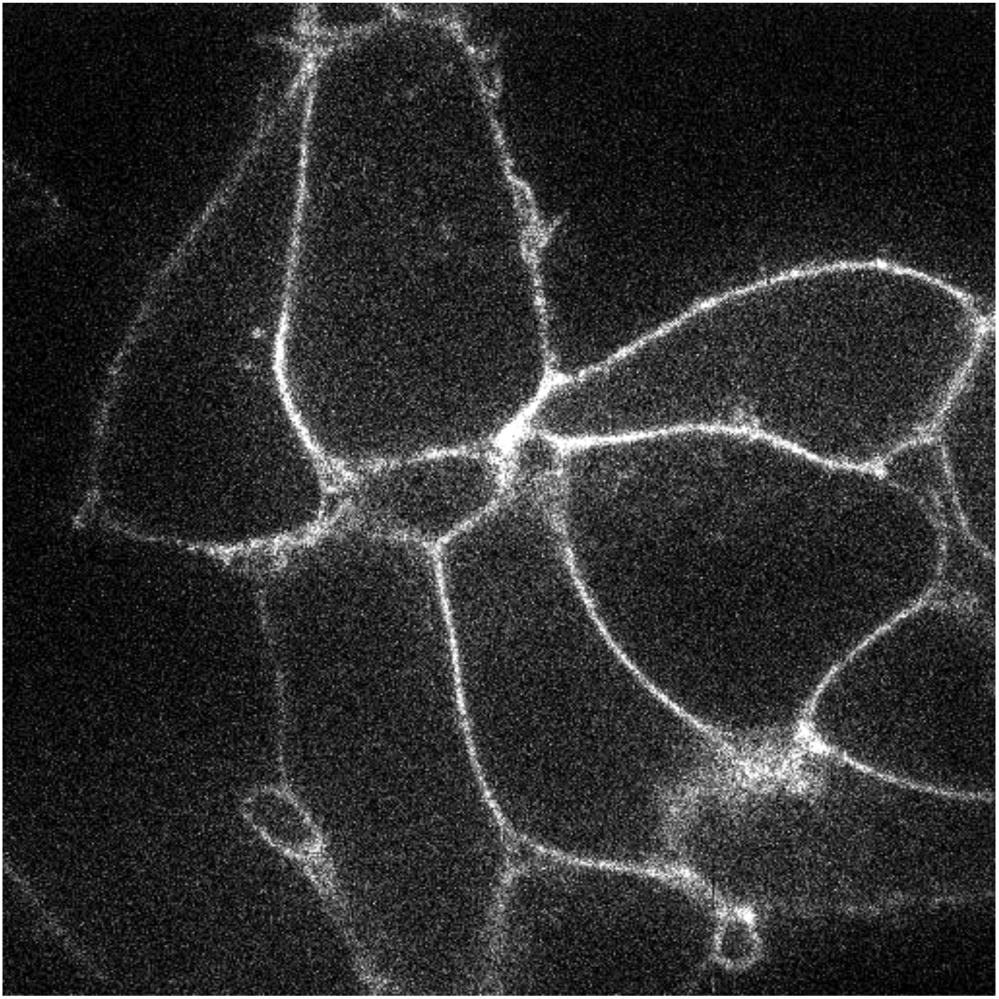
This video shows a confocal image series of AC9-EGFP overexpressed in a HEK293 cell. Cell was treated with 10µM isoproterenol added at t=0 in the time stamp. AC9 recruitment to internal puncta is observed over the course of 30 min.

**Supplementary Video 2.**
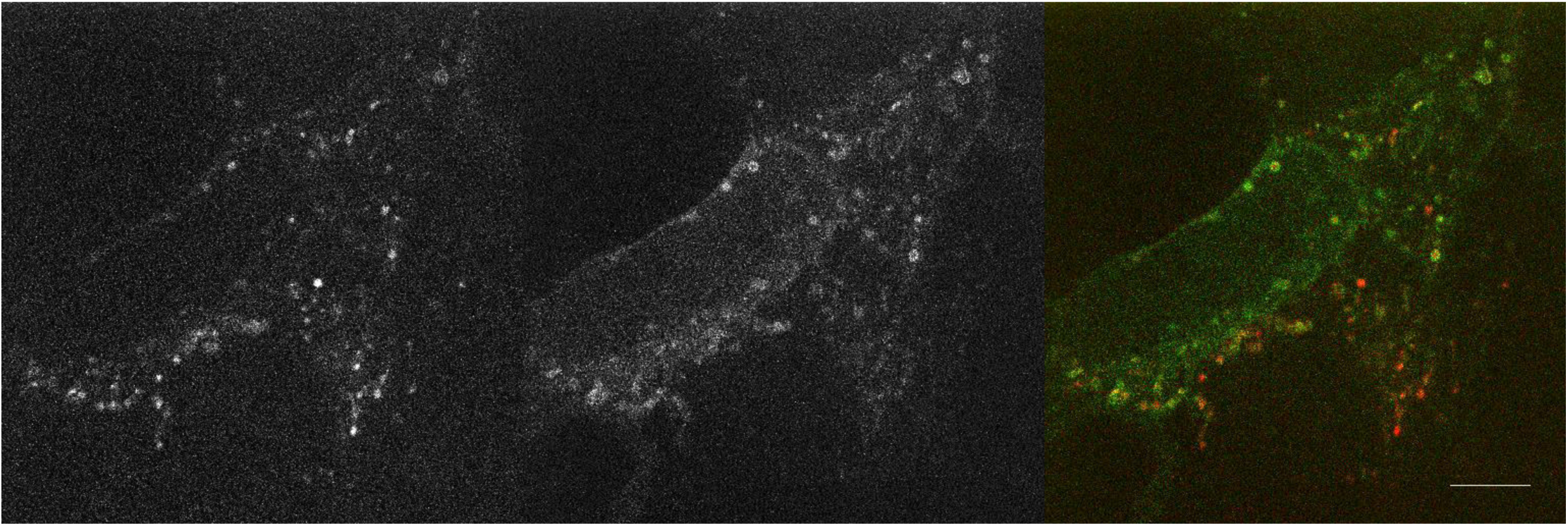
This video shows a confocal image series of β2AR (red) and Nb80-EGFP (green) from a HEK293 cell pre-incubated for 20 min with 100nM isoproterenol. 10µM alprenolol, added at t=0 time point indicated in the time stamp, reverses Nb80-EGFP recruitment to β2ARs at both the plasma membrane and endosomes.

**Supplementary Video 3.**
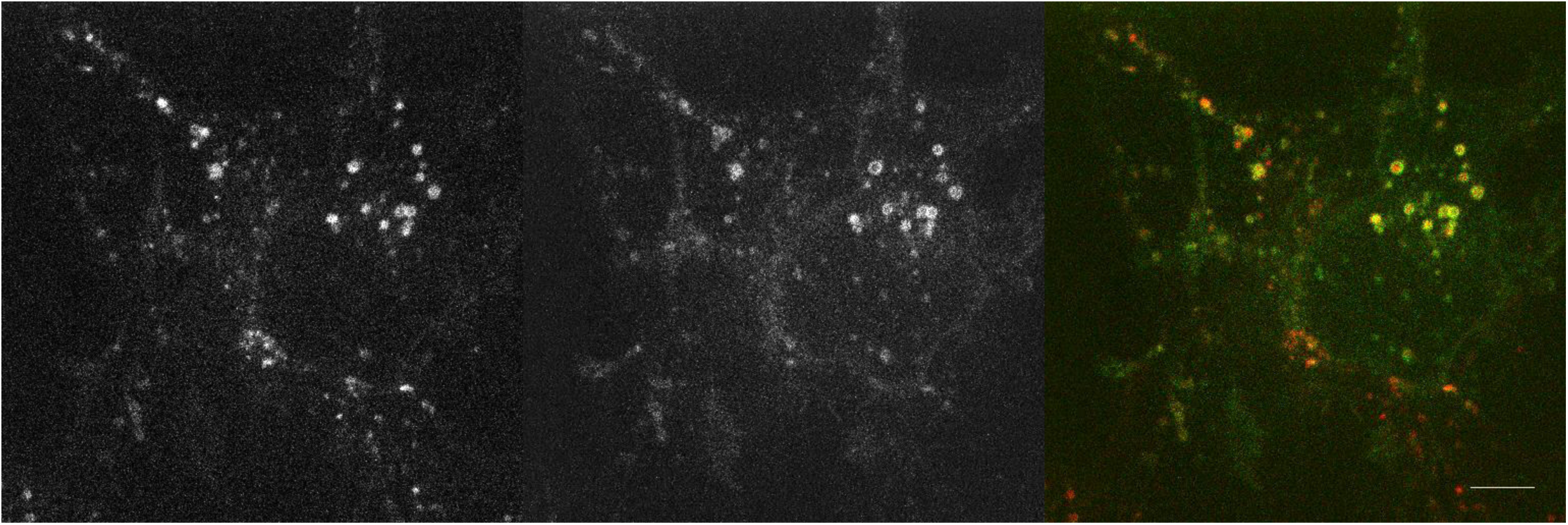
This video shows a confocal image series of β2AR (red) and Nb80-EGFP (green) from a HEK293 cell pre-incubated for 20 min with 100nM isoproterenol. 10µM CGP12177, added at t=0 indicated in the time stamp, reverses Nb80-EGFP recruitment to β2ARs at the plasma membrane but not at endosomes.

**Supplementary Video 4.**
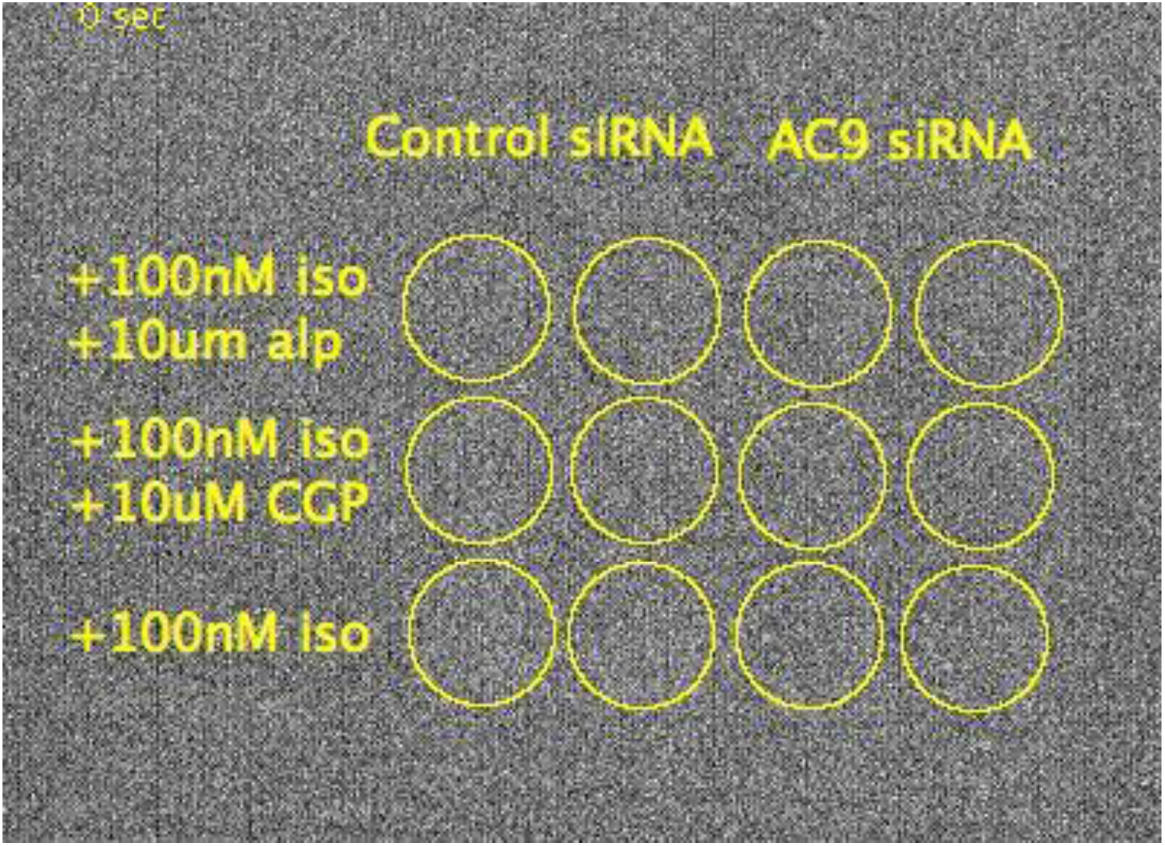
This video shows an image series of the luminescence-based cAMP biosensor from HEK293 cells that have been treated with control siRNA (columns 1 and 2) or AC9 siRNA (columns 3 and 4). Cells were preincubated with 100nM isoproterenol for 20 minutes prior to imaging, and 10µM CGP12177 or 10µM alprenolol were added immediately before imaging, where indicated.

## Materials and methods

### Cell culture, expression constructs, and transfections

HEK 293 cells (ATCC) were cultured in complete growth Dulbecco’s modified Eagle’s medium (DMEM, Gibco) and supplemented with 10% fetal bovine serum (UCSF Cell Culture Facility). HA-β2AR(Tang et al., 1999; von Zastrow and Kobilka, 1992), HA-V2R (Rochdi et al., 2010), HA-MOR (Whistler and von Zastrow, 1998), HA-V2R-T (Charest and Bouvier, 2003; Rochdi et al., 2010), all described previously, were sub-cloned from Flag-tagged constructs. Nb80-EGFP was previously described (Irannejad et al., 2013). HA-G(alpha)s, G(beta-1), G(gamma-2) were gifts from Philip Wedegaertner. HA-G(alpha)s-Q227L, a previously described point mutant of Gs that is constitutively active (Masters et al., 1989), was made from the original construct using the QuikChange Site-Directed Mutagenesis Kit (Agilent Technologies) with the forward primer 5’-CGATGTGGGCGGCCTGCGCGATGAACGCCGC-3’. Flag-AC1, Flag-AC9 from the Dessauer Lab, were originally described by (Hacker et al., 1998; Krupinski et al., 1989; Paterson et al., 2000; Premont et al., 1996). Flag-AC9-D442A (Catalytic inactive mutant) was also made from the original construct using QuikChange Kit with the forward primer 5’-CCACTAGTCCAGTGTGGTGGAATTCGCCATGGACTACAAAGACGATGACGAC-3’. Transfections were carried out using Lipofectamine 2000 (Life Technologies) according to the manufacturer’s protocol. Cells were transfected 48 hours before experiments. siRNA knockdown of AC1 and AC9 expression in HEK293 cells was carried out using Lipofectamine RNAiMAX (Life Technologies) according to the manufacturer’s protocol. Cells were transfected 72 hours before experiments. Knockdown of AC1 used the siRNA CCGGGCGGTTCAGACCTTCAA and AC9 knockdown used CTGGGCATGAGGAGGTTTAAA.

Primary cultures of human airway smooth muscle cells were established as described previously (Tsvetanova et al., 2016). Cells were passaged no more than 5 times using Trypsin-EDTA (Life Technologies) and maintained in 10% FBS in DMEM.

Gs knockout (Stallaert et al., 2017) and beta-arrestin-1/2 double knockout (O’Hayre et al., 2017) HEK293 cells were previously described. AC3 / AC6 double knockout HEK293 cells were also described previously (Soto-Velasquez et al., 2018) and were provided as a generous gift by Drs. Monica Soto-Valasquez and Val Watts (Purdue University). Cells were passaged using PBS-EDTA and maintained in 10% FBS in DMEM.

Cholera Toxin (Sigma) was administered to cells for 16 hours overnight treatment at 10 ng/ml concentration in 10% FBS in DMEM.

We found AC9 trafficking to be environmentally sensitive. Specifically, exposure of cells outside of the incubator for more than 2 minutes tended to reduce the degree of isoproterenol-stimulated internalization of AC9, without affecting internalization of β2AR. Accordingly, this restriction was consistently adhered to in the present study.

### Antibodies

Antibodies used were rabbit anti-Flag (Sigma), mouse anti-Flag M1 (Sigma), mouse anti-Flag M2 (Sigma), mouse anti-HA 16B12 (Biolegend), rat anti-HA (Roche), goat anti-AC9 (Santa Cruz Biotech), mouse anti-Golgin-97 (Thermo), rabbit anti-calnexin (Cell Signaling), mouse anti-Sodium/Potassium ATPase (Fisher).

### Fixed cell confocal imaging

Cells were transfected with the indicated construct(s) and then plated on glass coverslips coated with poly-L-lysine (0.0001%, Sigma) 24 hours later. For antibody feeding assays, cells were: (1) placed on ice and rinsed with ice-cold phosphate-buffered saline (PBS), (2) labeled by the addition of antibodies diluted 1:1000 in DMEM for 10 minutes, and (3) rinsed with room temperature PBS and allowed to traffic for 30 minutes by the addition of 37**°**C fresh media (DMEM + 10% fetal bovine serum) with or without a saturating concentration of β2AR agonist (10 µM isoproterenol, Sigma), V2R agonist (10 µM arginine-vasopressin, Sigma), MOR agonist (10 µM DAMGO, Sigma), or forskolin (10 µM, Sigma). For all assays, cells were rinsed with cold PBS and fixed by incubation in 3.7% formaldehyde (Fisher Scientific) diluted in modified BRB80 buffer (80mM PIPES, 1mM MgCl2, 1mM CaCl2, pH 6.8) for 20 minutes at room temperature. Cells were then blocked in 2% Bovine Serum Albumin (Sigma) in PBS with permeabilization by 0.2% triton X-100 (Sigma). Primary antibody labeling was performed by the addition of antibodies diluted 1:1000 in blocking/ permeabilization buffer for one hour at room temperature. Secondary labeling was performed by addition of the following antibodies diluted at 1:500 in blocking/ permeabilization buffer for 20 minutes at room temperature: Alexa Fluor 555 or 488 donkey anti-mouse (Invitrogen), Alexa Fluor 555 or 488 donkey anti-rabbit (Invitrogen), Alexa Fluor 488 or 555 goat anti-rat (Invitrogen), or Alexa Fluor 488 donkey anti-sheep (Life Technologies). Specimens were mounted using ProLong Gold antifade reagent (Life Technologies).

Fixed cells were imaged by spinning disc confocal microscope (Nikon TE-2000 with Yokogawa confocal scanner unit CSU22) using a 100X NA 1.45 objective. A 488 nm argon laser and a 568 nm argon/krypton laser (Melles Griot) were used as light sources.

### Microscope image acquisition and image analysis

Spinning disc images were collected using an electron multiplying CCD camera (Andor iXon 897) operated in the linear range controlled by Micro-Manager software (https://www.micro-manager.org). Images were processed at full bit depth for all analysis and rendered for display by converting to RGB format using ImageJ software (http://imagej.nih.gov/ij) and linear look up table. The number of endosomes was quantified by thresholding images and the ParticleTracker ImageJ plugin.

### Live-cell confocal imaging

Live cell imaging was carried out using Yokagawa CSU22 spinning disk confocal microscope with a x100, 1.4 numerical aperture, oil objective and a CO2 and 37 °C temperature-controlled incubator. A 488 nm argon laser and a 568 nm argon/krypton laser (Melles Griot) were used as light sources for imaging EGFP and Flag signals, respectively. Cells expressing both Flag-tagged receptor and the indicated nanobody-EGFP were plated onto glass coverslips. Receptors were surface labelled by addition of M1 anti-Flag antibody (1:1000, Sigma) conjugated to Alexa 555 (A10470, Invitrogen) to the media for 30 min, as described previously. Indicated agonist (isoprenaline, Sigma) or antagonist (CGP-12177, Tocris) (alprenolol, Sigma) were added and cells were imaged every 3 s for 20 min in DMEM without phenol red supplemented with 30 mM HEPES, pH 7.4 (UCSF Cell Culture Facility). Time-lapse images were acquired with a Cascade II EM charge-coupled-device (CCD) camera (Photometrics) driven by Micro-Manager 1.4 (http://www.micro-manager.org).

### Endosome Immunoisolation

Cells were transfected with the indicated construct(s) 48 hours before lysis and plated onto 60mm cell culture dishes 24 hours before lysis. Cells were allowed to traffic for 30 minutes by the addition of 37**°**C fresh media (DMEM + 10% fetal bovine serum) with or without a saturating concentration of the indicated agonist. Cells were then placed on ice, washed with ice-cold PBS, and scraped into an isotonic homogenization buffer (10mM HEPES, 100mM KCl, 25mM sucrose, Complete protease inhibitor (Roche), pH 7.2) and passaged 20 times through a 22 G BD PrecisionGlide Needle. Whole cell lysates were then spun down at 1000 G for 10 minutes at 4°C and the pellets discarded. The supernatant was then bound to Early Endosome Antigen 1 mouse antibody (1:250, Fisher Scientific) and anti-mouse IgG magnetic microbeads (Miltenyi Biotech) overnight. Endosomes were then bound to magnetic columns which were blocked with 3% BSA and washed with PBS. Proteins in the isolated fraction were eluted with 0.1% Triton-X and characterized by western blot.

### Surface Biotinylation

Cells were transfected with the indicated construct(s) 48 hours before lysis and plated onto 60mm cell culture dishes coated with poly-L-lysine (0.0001%, Sigma) 24 hours before lysis. Cells were allowed to traffic for 30 minutes by the addition of 37**°**C fresh media (DMEM + 10% fetal bovine serum) with or without a saturating concentration of the indicated agonist. Cells were then placed on ice, washed with ice-cold PBS, and then surface labeled with EZ-link Sulfo-NHS-biotin (Pierce) for 30 min, rocking at 4°C. Reaction was then quenched with tris buffered saline (TBS) twice for 10 min. Cells were then placed on ice, washed with ice-cold PBS, and scraped into an isotonic homogenization buffer (10mM HEPES, 100mM KCl, 25mM sucrose, Complete protease inhibitor (Roche), pH 7.2) and passaged 20 times through a 22 G BD PrecisionGlide Needle. Cell lysate was then bound to streptavidin agarose resin (Thermo) overnight. Resin was spun down and the supernatant discarded, resuspended and washed in ice-cold PBS, and characterized by western blot.

### Real-time cAMP assay in living cells

Real-time analysis of cAMP elevations were carried out in living HEK293 cells and in the absence of phosphodiesterase inhibitors using a were transfected with a plasmid encoding a cyclic-permuted luciferase reporter construct, based on a mutated RIIB cAMP-binding domain from PKA (pGloSensor-20F, Promega), which produces rapid and reversible cAMP-dependent activation of luciferase activity in intact cells and is capable of detecting cAMP elevations in the absence of phosphodiesterase inhibitors. Cells were plated in 24-well dishes containing approximately 200,000 cells per well in 500 µl DMEM without phenol red and no serum and equilibrated to 37 °C in a light-proof cabinet. An image of the plate was focused on a 512 x 512 pixel electron multiplying CCD sensor (Hamamatsu C9100-13), cells were equilibrated for 1 h in the presence of 250 µg ml^-1^ luciferin (Biogold), and sequential luminescence images were collected every 10 s to obtain basal luminescence values. The camera shutter was closed, the cabinet opened and the indicated concentration of isoprenaline was bath applied, with gentle manual rocking before replacing in the dark cabinet and resuming luminescence image acquisition. In endocytic manipulation experiments, cells were pre-incubated with 30 µM Dyngo-4a (abcam Biochemicals) for 15 min. Every 10 s, sequential images were acquired using Micro-Manager (http://www.micro-manager.org) and integrated luminescence intensity detected from each well was calculated after background subtraction and correction for vignetting using scripts written in MATLAB (MathWorks). In each multiwell plate, and for each experimental condition, a reference value of luminescence was measured in the presence of 5 µM forskolin, a manipulation that stimulates a moderate amount of receptor-independent activation of adenylyl cyclase. The average luminescence value-measured across duplicate wells-was normalized to the maximum luminescence value measured in the presence of 5 µM forskolin.

### Biochemical assay of cAMP accumulation

A biochemical assay of cAMP accumulation was used to determine the effects of AC mutation on catalytic activity, with high sensitivity and without dependence on subcellular location due to inhibition of cellular phosphodiesterases. Briefly, cells were pre-incubated in the presence of 1 mM IBMX (Sigma) for 30 min at 37°C in Dulbecco’s modified Eagle’s medium followed, and then incubated for an additional 10 min in absence or presence of isoproterenol (in the continued presence of IBMX), as indicated. Cells were quickly washed with ice-cold PBS and lysed by exposure to 0.1 M HCl for 10 minutes at room temperature. The cAMP concentration in lysates was determined using a commercial immunoassay (Direct cAMP ELISA kit, Enzo Life Sciences, Farmingdale, NY) according to the manufacturer’s instructions.

### Statistical Analysis

Results are displayed as the mean of results from each experiment or data set, as indicated in figure legends. The statistical significance between conditions for experiments with two conditions was calculated using paired, two tailed t-tests. All statistical calculations were performed using Excel (Microsoft Office) or Prism (GraphPad). The threshold for significance was p<0.05 and the coding for significance is reported as follows: (n.s.) p>0.05, (*) p:s0.05, (**) p:s0.01.

